# Transcriptional Elongation-Associated RNA Processing Errors in Induced Cellular Growth Arrest

**DOI:** 10.1101/2025.06.23.660200

**Authors:** Saeid Parast, Simai Wang, Marta Iwanaszko, Yue He, Deniz G. Olgun, Sarah R. Gold, Yuki Aoi, Jacob M. Zeidner, Benjamin C. Howard, William R. Thakur, Vijay Ramani, Ali Shilatifard

## Abstract

Transcription elongation factors control post-initiation steps of gene expression by RNA polymerase II (RNAPII). We have established distinct mechanistic roles for the essential elongation factors PAF1, NELF, SPT5, SPT6, and the Super Elongaiton Complex (SEC) via acute depletion of each individually in auxin-inducible degron lines. Here, we leverage these degron lines to explore the regulatory intersection of transcription elongation control and pre-mRNA processing. Integrating long- and short-read RNA-seq data to quantify transcript isoform usage at single-molecule resolution, we identify elongation factor-specific RNA processing regulons including a cellular senescence-enriched regulon shared by NELF and SPT6. We then show that long-term depletion of NELF or SPT6 results in reversible growth arrest following early upregulation of a small group of genes, which include the senescence-associated genes CDKN1A (p21) and CCN2. We perform genetic suppressor screens that implicate the elongation factor Elongin A (ELOA) in NELF or SPT6 depletion-induced growth arrest. ELOA loss suppresses NELF depletion-induced pre-mRNA processing defects and the 3’ extension of RNAPII occupancy past transcription end sites (TES) at genes induced by NELF depletion. ELOA also occupies TES-proximal regions under normal conditions, and acute ELOA depletion results in a loss of RNAPII processivity at the 3’ end of genes, opposing the effects of NELF or SPT6 depletion. Finally, we demonstrate that genetic loss of ELOA confers a growth advantage to aging human primary dermal fibroblasts. These findings establish the existence of novel ELOA-dependent mechanisms regulating transcription maturation, and links these mechanisms to the complex phenomena of cellular senescence and aging.

## Introduction

Transcription by RNA Polymerase II (RNAPII) is required for protein-coding gene expression in eukaryotes. The transcript elongation phase of this process, which is a critical step for the regulation of developmental and signal-responsive gene expression, is governed by a group of highly conserved elongation factors that regulate RNAPII pausing, elongation rate, processivity, and coordination with co-transcriptional processes such as pre-mRNA splicing and termination-coupled 3’-end processing (*1*). Key elongation factors include NELF (Negative Elongation Factor), DSIF, SPT6, and the PAF1 complex (PAF1C).

We have previously demonstrated that NELF is an essential factor required for normal transcriptional elongation genome-wide. In addition to its role in the post-initiation transcriptional rate-limiting step known as promoter-proximal pausing, NELF has also been linked to regulation of the rate of elongation by RNAPII and to the regulation of pre-mRNA processing steps associated with the RNAPII C-terminal domain (CTD) (5’ capping, splicing and termination-coupled 3’ transcript processing) (*2–5*). NELF and the pausing factor DSIF both bind to transcriptionally engaged RNAPII, causing it to arrest 20-60 nt downstream of the transcription start site (TSS) (*1*). The DSIF subunit SPT5 stabilizes paused RNAPII but also aids elongation by facilitating chromatin remodeling (*6–9*).

Paused RNAPII is released into processive elongation via the activity of the CDK9-containing factor positive transcription elongation factor b (P-TEFb), which phosphorylates NELF, DSIF, and Ser2 of the heptapeptide repeats within the RNAPII CTD, and phosphorylated NELF subsequently disassociates from RNAP (*10–13*). NELF was therefore thought to maintain pausing and repress elongation, as its name suggests (*8, 14*). However, we have previously demonstrated that NELF depletion is not sufficient and that NELF disassociation is not required for the release of paused RNAPII (*15*). Instead, NELF appears to be required for the first stage of a two-stage pausing process, and its disassociation from RNAPII may primarily allow for additional elongation factors to bind. We recently showed that SPT6 facilitates the displacement of NELF from promoter-proximally paused RNAPII by recruiting the PAF1 complex (PAF1C), enabling transition into productive elongation (*16*). SPT6 also facilitates efficient elongation and prevents premature termination via direct binding to the phosphorylated serine 2 and tyrosine 1 residues within the C-terminal domain (CTD) of RNAPII (*17, 18*). Finally, SPT6 ensures production of full-length transcripts by collaborating with factors like SPT5 and PAF1 to recruit elongation complexes and stabilize RNAPII on gene bodies (*18*).

Like NELF and SPT6, the RNAPII-associated factor ELOA has also been described as an elongation factor involved in elongation control. Along with ELOB and ELOC, ELOA was originally characterized as a key component of the transcriptional elongation factor complex Elongin, which was observed to enhance RNAPII processivity on naked DNA templates in cell-free in vitro systems (*19–21*). However, whether ELOA regulates RNAPII elongation in cells has been a matter of debate due to conflicting data indicating that ELOA deletion does not change the overall rate of elongation by RNAPII in vivo (*22, 23*). Instead, these in vivo studies showed that ELOA plays a role in the regulation of RNAPII fidelity in response to environmental stress and possibly developmental cues (*24, 25*). A more recent structure-based study shows that by binding to the RPB2 subunit of RNAPII through its “latch region” (residues 553-564), ELOA anchors the ELOB-ELOC heterodimer, which may indeed modulate RNAPII elongation rate (*26*). Via its Cullin-box sequence, ELOA (still bound to ELOB-ELOC) can also form an E3 ubiquitin ligase complex with CUL5/RBX2 (*27, 28*). Nevertheless, the exact nature of the role(s) ELOA plays in elongation control, its interplay with other elongation factors, and its function in the regulation of specific transcriptional programs in response to cellular stress conditions remain elusive.

Cellular senescence, a phenotypic response to stress conditions that is closely related to aging, is characterized as a state of permanent or irreversible cell cycle arrest in previously proliferating cells (distinct from terminal differentiation); this arrest occurs due to inhibition of the cyclin-dependent kinases (CDKs) CDK4/6 by p16/p21 (CDKN1A/CDKN2A), resulting in hypo-phosphorylation of retinoblastoma protein (Rb), E2F sequestration and subsequent downregulation of cell cycle progression genes (*29, 30*). It is widely believed that transcription by RNAPII is decreased in senescent cells, and more promoter-proximal pausing is observed (*31, 32*). Similarly, we have found that most but not all RNAPII-transcribed genes are paused upon depletion of NELF-C/D (NELF-C), a condition that also induces growth arrest (*15*). Senescent cells remain metabolically active and acquire a senescence-associated secretory phenotype (SASP), which includes the expression and secretion of various factors that are cell type and context-specific but generally include inflammatory factors such as IL-6 and growth factors such as CCN2 (*33–35*).

Here, we first investigate the effects of acute elongation factor depletion on transcript isform usage in degron cell lines using long-read Iso-seq data, which we quantify via integration with data from short-read RNA-seq. Interestingly, we find that loss of either NELF-C or SPT6 but neither SPT5 nor PAF1 results in a gene signature associated with cellular senescence. We then investigate the effects of NELF-C, NELF-E, SPT5 or SPT6 depletion on cell growth and identify a reversible growth arrest phenotype in cells depleted of NELF-C, NELF-E, or SPT6, but not SPT5 (for which depletion results in cell death). In auxin-treated NELF-C-AID cells, we observe upregulated expression of senescence-associated genes and we identify a set of 111 genes at which RNAPII is released upon NELF-C depletion rather than pausing at a second site. RNAPII is similarly released at these genes upon SPT6 depletion in SPT6-AID cells.

We perform a CRISPR screen for sgRNA KO-conferred rescue of NELF-C and SPT6 depletion-induced growth defects. Among the top hits we identify Elongin A (ELOA), a member of the Elongin complex with a long-disputed role in transcriptional elongation control. We then confirm that ELOA mediates the growth arrest induced by NELF or SPT6 depletion and show that its overexpression amplifies the NELF-C depletion-induced expression of senescence-associated genes. We also show that the primate-specific ELOA homolog ELOA3 can play a similar role. Subsequent long-read analysis of transcript isoforms upon NELF depletion by Iso-Seq reveals the emergence of fusion transcripts indicative of potential defects in transcription termination and an ELOA-dependent shift that is specific to genes where RNAPII is released in NELF-depleted cells. We demonstrate that the genes where RNAPII is released upon NELF depletion are ELOA-occupied genes, and that dramatically increased ELOA levels at ELOA-occupied genes underlies 3’ RNAPII occupancy past transcription end sites. Consistent with this, we report marked 3’ elongation defects upon acute ELOA depletion for a unique cluster of short genes at which ELOA accompanies elongating RNAPII. Finally, we demonstrate that ELOA confers a growth advantage to aging human fibroblasts.

## Results

### Single-molecule transcriptomic profiling reveals elongation factor specific RNA processing regulons

The elongation factors PAF1C, SPT5, SPT6, and NELF are essential for cell viability. Our group has previously engineered auxin-inducible degron (AID) cell lines to rapidly and efficiently deplete each of these essential elongation factors (*6, 7, 15, 16, 36*), allowing us to isolate the impact of each factor on the regulation of transcriptional and co-transcriptional processes. We sought to leverage these cell lines to explore how transcription elongation regulation intersects with RNA processing. Short-read RNA-seq methods deliver high read depth and are therefore powerful for quantifying gene expression, but they are limited for accurately quantifying distinct or novel transcript isoforms that reflect altered RNA processing decisions (*e.g.* transcription start and termination site usage; exon usage and intron retention). Long-read sequencing methods such as Iso-seq (PacBio) deliver nearly unambiguous correspondence between read and isoform and are particularly powerful for novel isoform discovery, but they lack read depth and introduce sequence and length biases. To account for these complementary strengths and limitations, we chose to jointly sequence steady-state polyadenylated transcriptomes in bulk and at single-molecule resolution, via short-read Illumina and long-read PacBio sequencing, respectively. We sequenced mRNA purified from NELF-C-AID, SPT5-AID, SPT6-AID, and PAF1-AID degron lines treated with auxin or DMSO for 6h, as well as parental OSTIR1-expressing DLD-1 cells (as a negative control). The 6h auxin treatment time was chosen to enable efficient elongation factor depletion while minimizing secondary effects. Following published data processing pipelines (*37, 38*) for both platforms, we then quantified transcript isoform usage using the MPAQT (*39*) algorithm (Fig. 1A). In total and across all cell lines, our dataset comprises 10.36M molecules mapping to 33,050 genes, with 132,095 isoforms falling within a wide variety of categories (Fig. 1B).)

**Figure 1.**
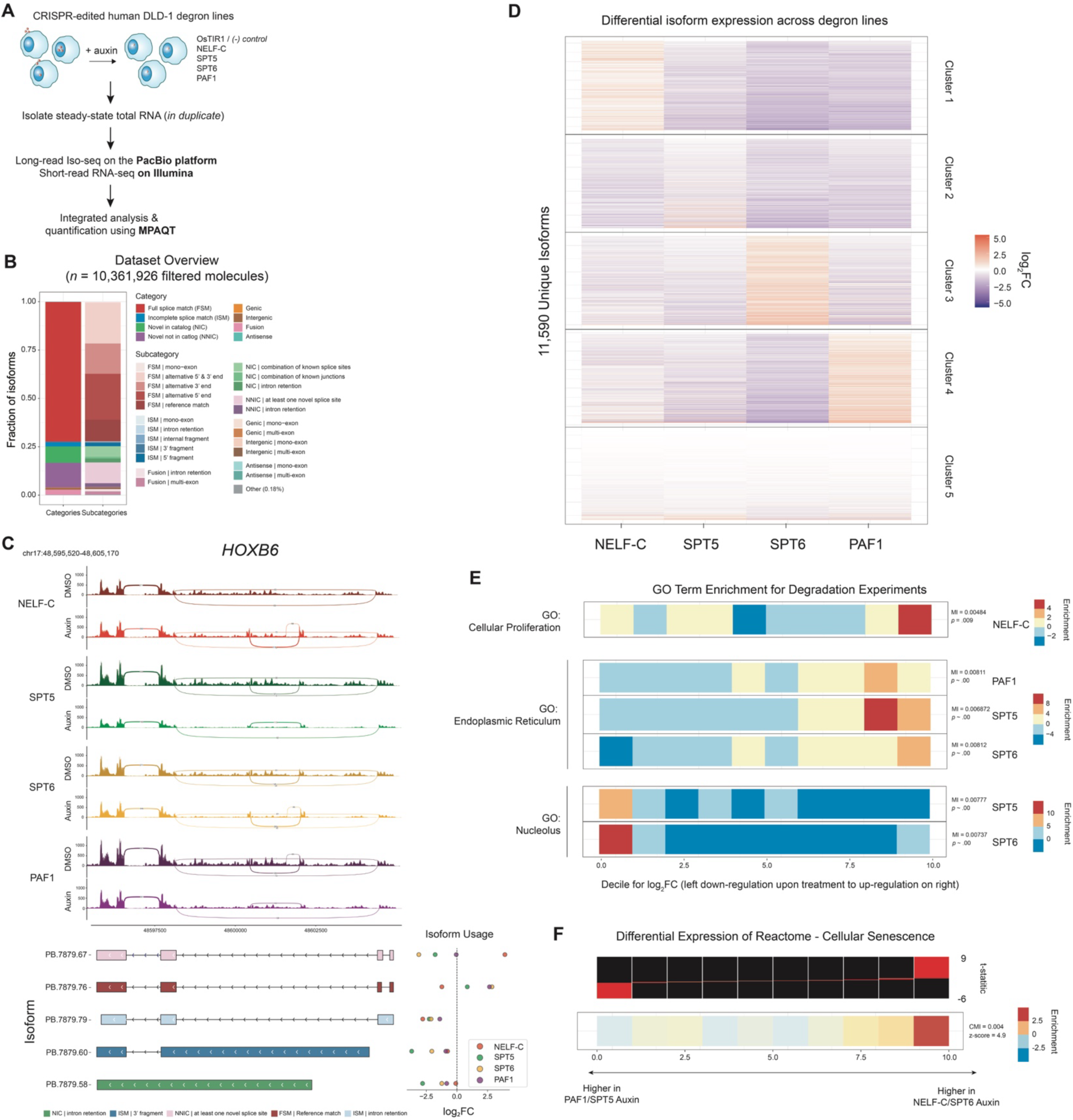
Iso-Seq-based identification and integrated quantification of novel transcripts in acute RNAPII elongation factor depletion. (A) Schematic overview of data generation and analysis in this transcriptomic profiling study. Acute depletion of the RNAPII elongation factors NELF-C, SPT5, SPT6, PAF1 was accomplished via auxin treatment in their respective DLD-1 degron cell lines; parental DLD-1 cells expressing the OsTIR1 E3 ligase were included as a negative control. Total RNA was isolated and prepared into libraries for long- and short-read sequencing. Long-read libraries (polyadenylated full-length transcripts) were sequenced on the PacBio platform using the Iso-seq method. Short-read libraries were sequenced on the Illumina platform. Isoform-resolved transcript quantification was conducted using MPAQT. (B) Dataset overview of the transcriptome across all acute depletion experiments, as defined by Iso-seq. Over 10 million full-length reads were tabulated. Isoform types were defined using SQANTI3, including full splice matches (FSM), incomplete splice matches (ISM), genic, intergenic, fusion, antisense transcripts, novel transcripts that use known splice sites / junctions (NIC) or novel splice sites / junctions (NNIC), and their respective subtypes. (C) Sashimi (*58*) plot of short-read RNA-seq data for comparative visualization of exon usage (top) and track visualization of isoforms identified by Iso-seq (bottom) at the example gene HOXB6, for each of the degron lines described in (A). The five most highly expressed isoforms isoforms (across all conditions) and their SQANTI3 subtypes are shown. Changes in isoform usage upon acute RNAPII elongation factor depletion are shown as log2 fold change along the isoform tracks. (D) K-means clustering of differential expression for 11,590 isoforms with |logFC| ≥ 1 in at least one of the factor knockdown contrasts. (E) GO term enrichment for genes associated with the up- and down-regulated isoforms in (D), indicating distinct functional consequences for depletion of different elongation factors. Enrichment is shown for the cellular proliferation GO term in NELF-C degron cells, for the ER regulation term in SPT5, SPT6 and PAF1 degron cells, and for the nucleolus term in SPT5 and SPT6 degron cells; enrichment is shown for each decile of log2FC upon auxin treatment. (F) Heatmap illustrating enrichment analysis for the REACTOME_CELLULAR_SENESCENCE gene set upon acute depletion of NELF-C or SPT6 relative to acute depletion of PAF1 or SPT5. The top panel displays the distribution of isoform t-statistics (limma-trend) averaged by gene across ten equally populated bins, with black bars representing the overall t-statistic range and red overlays delineating the bin-specific ranges. The bottom panel shows the enrichment scores for the gene set within each bin as calculated by pyPAGE. To the right, the Conditional Mutual Information (CMI) value calculated by pyPAGE quantifies the association between gene expression changes and gene set membership while accounting for annotation biases, with its z-score indicating the number of standard deviations by which the observed CMI deviates from the distribution of all CMI values derived from an expanded gene set list.

Our approach resulted in an isoform-resolved transcriptomic ‘atlas’ of distinct elongation factor perturbations. We note that steady-state measurement of mature RNA abundance necessarily underestimates the full effect of these perturbations, as it does not capture nascent RNA, including pre-mRNA processing intermediates. Nevertheless, the integration of short- and long-read data allows for quantitative assessment of differential isoform usage across all experimental conditions, as visualized for the *HOXB6* gene (Fig. 1C).

We used MPAQT-based, per-isoform log2 fold-change (log2FC) estimates to explore the consequences of each perturbation on differential isoform usage, employing k-means clustering (*k* = 5) across all samples for 11,590 filtered isoforms with |logFC| ≥ 1 in at least one of the factor knockdown contrasts. This clustering analysis (Fig. 1D) revealed distinct sets of upregulated isoforms. While a large set of isoforms (Cluster 5) change in a factor-independent manner, other isoforms (Clusters 1 - 4) appear to be upregulated in response to perturbation of specific elongation factor(s), implying that transcription elongation factors may have transcript-specific regulatory effects.

We next sought to address the functional significance of differentially upregulated isoforms for each elongation factor. We performed functional enrichment tests using the pyPAGE algorithm (*40*) to assess the statistically significant enrichment or depletion of up- and down-regulated isoforms. Intriguingly, we observe distinct functional enrichments associated with each perturbation (relative enrichment plots and associated statistical significance tabulated in Fig. 1E). For instance, genes associated with the GO term ‘cellular proliferation’ were significantly upregulated upon NELF depletion, with no significant associations observed for the same gene set across the other perturbations. Conversely, upregulated genes were significantly co-associated with the GO term ‘endoplasmic reticulum’ for depletion of SPT5, SPT6, or PAF1, suggesting some level of compartmentalization of regulation by these factors. Finally, downregulated genes were significantly co-associated with the GO term ‘nucleolus’ for depletion of SPT5 or SPT6, suggesting a possible association between rRNA processing and these two elongation factors that may be further explored in future studies. Taken together, these analyses suggest distinct transcriptomic and functional consequences of acute elongation factor depletion in DLD-1 cells.

### Long-term NELF-C or SPT6 depletion results in reversible cellular growth arrest

We previously reported that long-term depletion of NELF-C via auxin treatment in NELF-C AID cells results in a severe growth defect, with evidence of cell death beginning after 24h (*15*). To further investigate the effects of NELF subunit depletion, we initially performed a long-term growth assay in which either NELF-C, NELF-E, or SPT5 were depleted via addition of auxin for four consecutive days. As expected, growth in auxin-treated NELF-C, NELF-E, and SPT5 AID cells was severely impaired relative to DMSO-treated cells (Fig. 2A, Fig. S1A). However, we then observed that NELF-depleted cells could resume growth upon wash-off of the auxin-containing media (Fig. 2A, Fig. S1A), indicating that long-term NELF depletion results in growth arrest that eventually leads to cell death but is at least initially reversible. In contrast, SPT5-depleted cells did not resume growth upon auxin wash-off (Fig. 2A), indicating that long-term SPT5 depletion results in irreversible growth arrest and cell death. Next, we performed genome-wide studies to characterize early changes in RNAPII transcription associated with the NELF-C depletion-induced phenotype that we observed in the long-term growth assay. An auxin treatment time of 6 hours was chosen to ensure efficient NELF-C depletion while also minimizing indirect or secondary effects. Consistent with pathway enrichment analysis of the NELF-C-depleted transcriptome (Fig. 1F), RNA-seq analysis revealed a dramatic increase in the mRNA expression levels of the cellular senescence-associated genes *CCN2* and *CDKN1A* (p21) upon short-term NELF-C depletion (Fig. 2B-C). Upregulated gene expression is contrary to the genome-wide trend we previously reported, in which NELF-C depletion causes RNAPII to pause at a second pause site associated with the +1 nucleosome. However, consistent with our RNA-seq analysis, paired ChIP-seq analyses revealed a dramatic increase in the occupancy of RNAPII and its elongation-associated CTD modification Ser2P within the gene body at both the *CCN2* and *CDKN1A* loci, accompanied by the elongation factors SPT6 and PAF1 (Fig. 2B-C). ChIP-seq also confirmed that this transcriptional induction occurred despite efficient depletion of NELF (Fig. 2B-C). Together, these results indicate that short-term NELF-C depletion induces the release of RNAPII into productive elongation at these senescence-associated genes.

**Figure 2.**
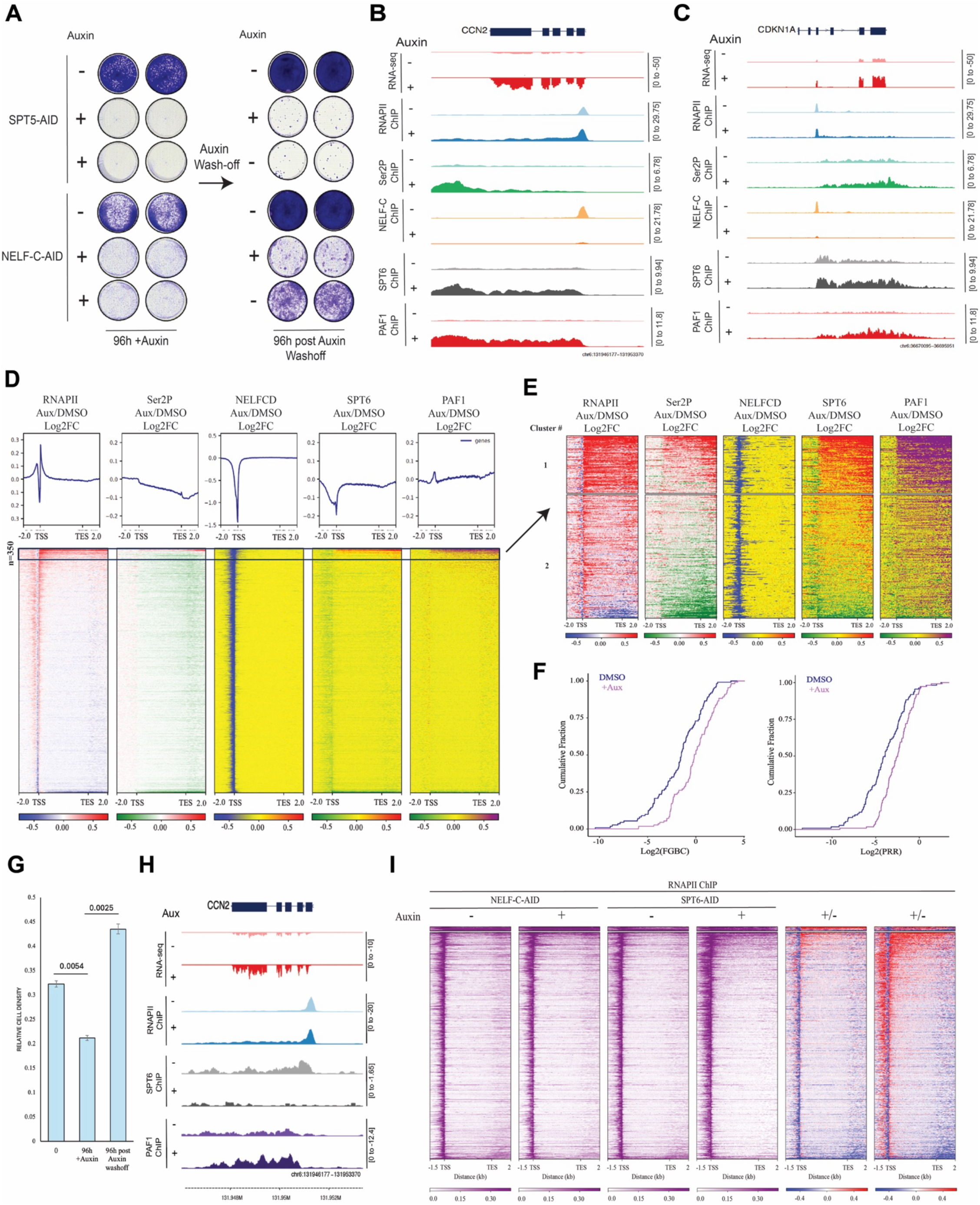
NELF depletion induces growth arrest concomitant to RNAPII release at senescence-associated genes. (A) Reversible growth defect observed upon auxin treatment in NELF-C-AID but not SPT5-AID cells. Cells were treated with auxin (Aux, +) or DMSO control (-) for 96h prior to wash-off and culture in fresh media for 96h, followed by fixation and crystal violet staining to visualize confluency. (B-C) Genome browser track visualization of RNA-seq and ChIP-seq signals at the senescence-associated genes CCN2 (B) and CDKN1A (C) in NELF-C-AID cells treated with DMSO or auxin for 6h. ChIP-seq was performed using antibodies against RNAPII, Ser2P (elongation-associated modification of the RNAPII CTD), NELF-C, SPT6 and PAF1. (D) Genome-wide metaplots of Log2-transformed fold change (Log2FC, above) and heatmaps (below) of differential ChIP-seq signal (RNAPII, Ser2P, NELF-C, SPT6, PAF1) in auxin-treated relative to DMSO-treated NELF-C-AID cells. TSS= transcription start site; TES=transcription end site. n=6038 genes. In the differential ChIP-seq signal heatmaps, the 350 genes with the strongest differential RNAPII signal within the gene body are highlighted. (E) Expanded view of the differential ChIP-seq signal heatmaps for the 350 genes highlighted in (D), divided further into two clusters based on strength of differential RNAPII signal in the gene body. Upper cluster (cluster 1), n=111 genes; lower cluster, n=239 genes. (F) Empirical distribution functions (ECDF) plotted for Log2-transformed full gene body coverage ratio (Log2(FGBC), left) and Log2-transformed pause release ratio (Log2(PRR), right) calculated from RNAPII ChIP-seq reads at cluster 1 genes (n=111) in auxin-treated cells (pink) versus DMSO-treated cells (blue). (G) Bar graph showing relative cell density in SPT6-AID cells treated with auxin for 96h prior to wash-off and culture in fresh media for 96h. (H) Genome browser track visualization of ChIP-seq signals at the senescence-associated gene CCN2 in SPT6-AID cells treated with DMSO or auxin for 6h. ChIP-seq was performed using antibodies against RNAPII, Ser2P (elongation-associated modification of the RNAPII CTD), SPT6 and PAF1. (I) Genome-wide metaplots of Log2-transformed fold change (Log2FC, above) and heatmaps (below) of differential RNAPII ChIP-seq signal in auxin-treated relative to DMSO-treated NELF-C-AID and SPT6-AID cells. TSS= transcription start site; TES=transcription end site. Clustering is applied from Fig. 2 D-E; n=6433 genes.

Global analysis of our ChIP-seq data revealed that among all RNAPII-occupied genes (n=6038), a very small cluster (cluster 1, n=111) showed robust release of RNAPII into the gene body upon NELF-C depletion (Fig. 2D-E, Fig. S1B-C). Cluster 1 genes also showed significant increases in NELF-C depletion-induced SPT6 and PAF1 occupancy relative to other RNAPII-occupied genes (Fig. 2E, Fig. S1B-C). The impact of NELF-C depletion on the transcription of genes in cluster 1 is particularly striking when RNAPII release is quantified as pause release ratio (PRR) and processive elongation is quantified as full gene body coverage ratio (FGBC) (Fig. 1F). We also performed genome-wide studies in SPT6-depleted cells, which show a similarly reversible growth arrest phenotype (Fig. 2G). RNA-seq analysis revealed induction of *CCN2* expression upon short-term SPT6 depletion (Fig. 2H). Consistent with this, paired ChIP-seq analyses revealed clear increases in RNAPII and PAF1 occupancy within the gene body at the *CCN2* locus upon short-term SPT6 depletion (Fig. 2H). Side by side comparison of global RNAPII occupany profiles via ChIP-seq in NELF-C and SPT6-AID cells revealed a similar activation of cluster 1 genes upon short-term depletion of NELF-C or SPT6, corroborating the results of our transcriptomic analysis, which indicated the activation of similar gene signatures upon NELF-C or SPT6 depletion (Fig. 2I). Together, these results indicate that early upregulation of a small group of genes (which includes CCN2, CDKN1A, and other senescence-associated genes) plays a role in the reversible growth arrest phenotypes induced by long-term NELF or SPT6 depletion.

### Suppressor genetic screen identifies ELOA mediating NELF-C or SPT6 depletion-induced cellular growth arrest

To investigate the molecular mechanisms by which long-term NELF-C depletion can induce a growth arrest phenotype, we performed a genome-wide CRISPR-Cas9 screen for resistance to NELF-C depletion-induced growth impairment upon auxin treatment in the NELF-C-AID DLD-1 cell line (Fig. 3A). Comparison of auxin-treated NELF-C-AID to auxin-treated parental cells (Fig. S2A) revealed a strong enrichment of sgRNAs targeting ELOA (labelled as TCEB3), with weaker enrichment of sgRNAs targeting ELOB (TCEB2) and ELOC (TCEB1), in NELF-C-AID cells compared to parental cells (Fig. 3B, Fig. S2B). To validate the CRISPR screen results, we first infected NELF-C-AID DLD-1 cells with two independent sgRNAs targeting ELOA to generate distinct, clonal ELOA KO cell lines (Fig. S2C-F). Next, we used these clonal ELOA KO cell lines to assess the impact of ELOA KO on the growth arrest phenotype previously observed upon NELF-C depletion. ELOA KO appeared to abrogate the growth arrest in auxin-treated NELF-C-AID cells (Fig. 3C). RNA-seq analysis revealed that ELOA KO also diminished the NELF-C depletion-induced upregulation of CCN2 and CDKN1A (Fig. 3D-E). This impact was recapitulated at the protein level, where ELOA KO also abrogated NELF-C depletion-induced PARP cleavage (Fig. 3F). ELOA KO similarly diminished the NELF-C depletion-induced upregulation of cluster 1 genes (Fig. S3B), consistent with a putative role for ELOA as an elongation factor. ELOA KO also diminished the NELF-C depletion-induced expression of a larger gene set (n=455) (Fig. 3G). In fact, ELOA KO alters the expression of many genes (n=551) irrespective of the presence or relative absence of NELF-C (Fig. S3C-D), indicating a broader role for ELOA as a transcriptional regulatory factor beyond the artificial condition of auxin-induced NELF-C depletion.

**Figure 3.**
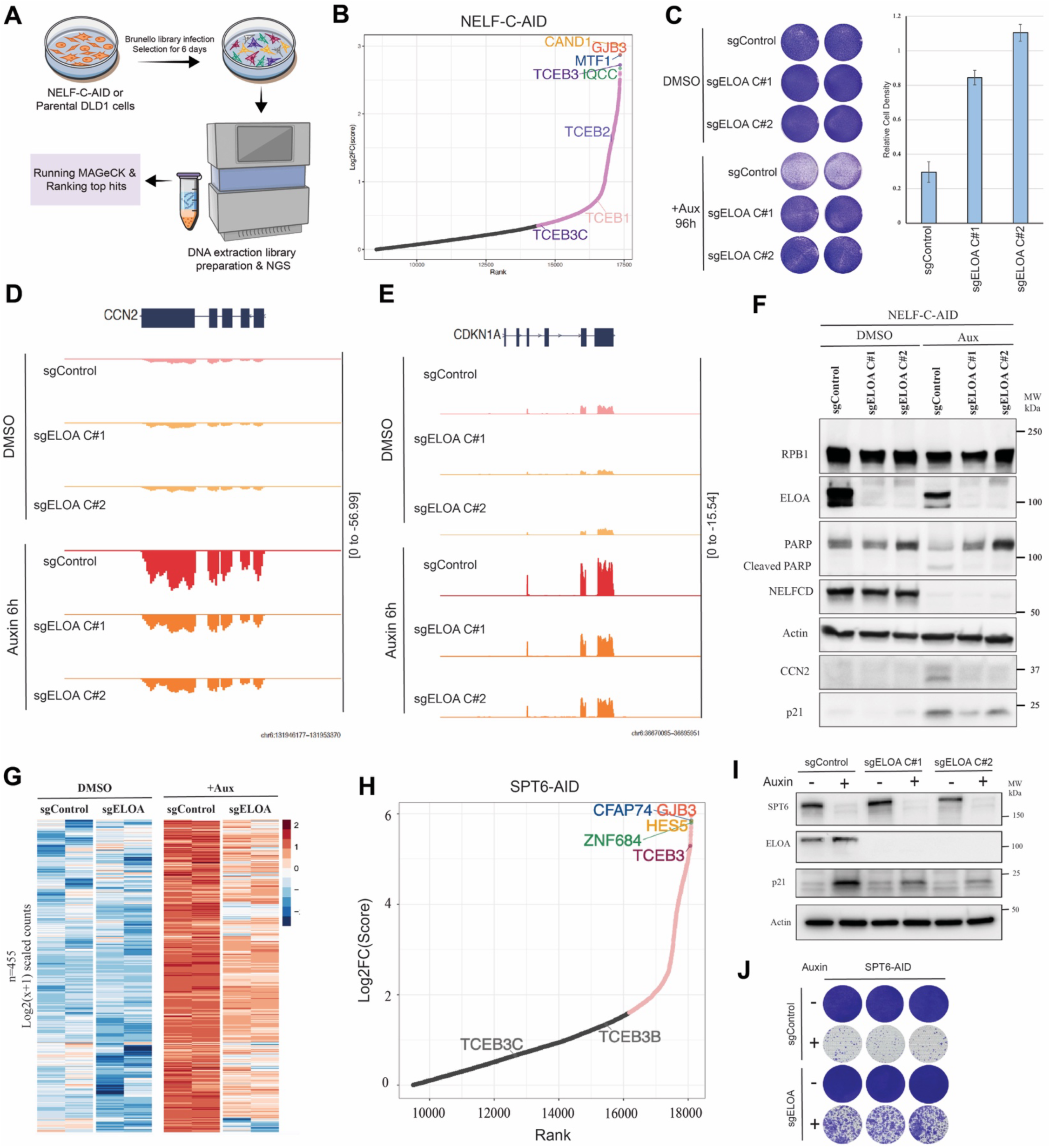
ELOA knockout abrogates the phenotype similarly induced by NELF-C or SPT6 depletion. (A) Schematic illustration of the Brunello library CRISPR screen to identify factors limiting the growth of NELF-C-AID cells upon auxin-induced NELF depletion. Parental (OSTIR1-expressing) DLD-1 cells were used as a control for the effects of auxin treatment. (B) Rank plot of Log2-transformed fold change (Log2FC) in gene essentiality score (β-score), the output from MAGeCK-mle analysis of the CRISPR screen in NELF-C-AID DLD-1 cells. Top hits including ELOA (TCEB3) are highlighted, as are the Elongin family members ELOB (TCEB2), ELOC (TCEB1) and ELOA3 (TCEB3C). (C) Rescue of NELF depletion-induced growth defect in NELF-C-AID cells via clonal knockout of ELOA using two distinct ELOA-targeting sgRNA (ELOA-KO-C#1 and #2), with a nontargeting sgRNA as control (sgControl). Cells were treated with auxin (+Aux) or DMSO control (-Aux) for 96h. The number of auxin-treated relative to DMSO-treated cells is quantified at right. (D-E) Genome browser track visualization of RNA-seq signal at the senescence-associated genes CCN2 (D) and CDKN1A (E) in sgControl and sgELOA (ELOA KO 1 and 2) NELF-C-AID cells treated with DMSO or auxin. (F) Western blots for the RNAPII subunit RPB1, for ELOA, for PARP and its cleavage product, for NELFCD to confirm depletion, for actin (as a loading control), and for the senescence-associated proteins CCN2 and p21, in NELF-C-AID cells treated with DMSO or auxin for 24h. (G) Heatmap of gene expression levels (Log2-transformed, pseudocount-adjusted scaled RNA-seq read counts) in sgControl versus sgELOA NELF-C-AID cells treated with DMSO or auxin for 6h, for the set of genes induced upon auxin treatment in sgControl cells (n=455). (H) Rank plot of Log2-transformed fold change (Log2FC) in gene essentiality score (β-score), the output from MAGeCK-mle analysis of the CRISPR screen in SPT6-AID DLD-1 cells. Top hits including ELOA (TCEB3) are highlighted. (I) Western blots for SPT6 to confirm depletion, for ELOA to confirm knockout, for actin (as a loading control), and for the senescence-associated protein p21, in SPT6-AID cells treated with DMSO or auxin for 24h. (J) Rescue of SPT6 depletion-induced growth defect in SPT6-AID cells via clonal knockout of ELOA using two distinct ELOA-targeting sgRNA (ELOA-KO-C#1 and #2), with a nontargeting sgRNA as control (sgControl). Cells were treated with auxin (+Aux) or DMSO control (-Aux) for 96h.

To further validate ELOA as a mediator of senescence-associated gene expression and growth arrest resulting from NELF depletion, we first generated a NELF-C-dTAG line in the human SYO-1 cell line background (Fig. S4A), which is characterized as having a low mutational burden (*41*). The effect of dTAG13 treatment-induced NELF-C depletion in these NELF-C-dTAG cells was similar to that of auxin-induced NELF-C depletion in NELF-C-AID DLD-1 cells: NELF-C depleted cells stop growing but do not die, and they begin to proliferate again upon dTAG-13 wash-off (Fig. S4B). Next, we generated a NELF-C and ELOA double dTAG cell line in the same SYO-1 background. We were able to efficiently deplete both NELF-C and ELOA proteins via addition of dTAG-13 into the culture medium (Fig. S4C). Consistent with results of ELOA KO in auxin-treated NELF-C-AID DLD-1 cells, the NELF-C depletion-induced growth arrest observed in dTAG13-treated NELF-C-dTAG SYO-1 cells could be partially rescued via the additional depletion of ELOA in this double dTAG SYO-1 cell line (Fig. S4D).

Finally, because we observed a similar transcriptional disruption upon loss of either NELF or SPT6 (Fig. 1F, Fig. 2I), we hypothesized that the growth arrest observed in auxin-treated SPT6-AID cells could also be mediated by ELOA, similar to auxin-treated NELF-C-AID cells. To investigate this, we performed a similar unbiased genome-wide CRISPR screen for resistance to SPT6 depletion-induced growth impairment upon auxin treatment in the SPT6-AID DLD-1 cell line (Fig. 3H). As expected, ELOA (labelled as TCEB3) was a top hit of this screen (Fig. 3H). Next, we generated distinct, clonal ELOA KO cell lines by infecting SPT6-AID DLD1 cells with two independent sgRNAs targeting ELOA (Fig. 3I). ELOA KO abrogated the SPT6 depletion-induced increase in CDKN1A protein level (Fig. 3I). ELOA KO also appeared to abrogate the growth arrest in auxin-treated SPT6-AID cells (Fig. 3J). Together, these results point to ELOA as a transcriptional regulatory factor that mediates the reversible growth arrest phenotype observed upon NELF-C or SPT6 depletion, including the NELF-C or SPT6 depletion-induced upregulation of senescence-associated genes and protein markers.

### ELOA overexpression amplifies the impacts of NELF-C depletion

To further substantiate ELOA as an elongation factor that mediates the reversible growth arrest phenotype induced by NELF-C depletion, we used a tetracycline (Tet)-inducible system to stably overexpress an N-terminal FLAG- and HA-tagged ELOA construct upon treatment with doxycycline (Dox) in our NELF-C-AID cells (Fig. 4A, Fig. S5A). Dox-induced overexpression of ELOA (but not GFP) enhanced senescence-associated β-galactosidase activity in auxin-treated NELF-C-AID cells (Fig. 4B). Dox-induced overexpression of ELOA also amplified upregulation of the senescence-associated genes CCN2 and CDKN1A upon NELF-C depletion but had no effect on the expression of these genes in the presence of NELF-C (Fig. 4C-D). Cluster 1 genes (at which RNAPII is released upon NELF-C depletion) demonstrated a similar trend of ELOA-amplified upregulation in response to NELF-C depletion, with no clear effect of ELOA overexpression on cluster 1 genes in the presence of normal NELF-C levels (Fig. 4E, S5B-D). Consistent with these findings, ChIP-seq for NELF and the HA tag revealed that while HA-ELOA overexpression does not appreciably impact NELF occupancy at CCN2, ID2, or other cluster 1 genes, NELF-C depletion caused a strong increase in the occupancy of HA-ELOA throughout the body of these genes (Fig. 4F-H, Fig. S5E). We also performed ChIP-seq for RNAPII and the elongation factors SPT6 and PAF1, which have been shown to associate with ELOA (*22, 42*). In both the presence and relative absence of NELF, Dox-induced overexpression of ELOA had no appreciable effect on RNAPII occupancy at cluster 1 genes (Fig. S5D). However, ELOA overexpression amplified the NELF-C depletion-induced occupancy of SPT6 throughout the body of cluster 1 genes, in close association with the pattern of NELF-C depletion-induced ELOA occupancy detected by HA-tag ChIP-seq (Fig. 4H, Fig. S5E). Together, these results indicate that the loss of NELF-C from the promoters of senescence-associated genes either allows for or causes (e.g. via transcriptional stress) ELOA to associate with SPT6-equipped RNAPII within the gene body, leading to increased expression of this set of genes and the consequent senescence phenotype.

**Figure 4.**
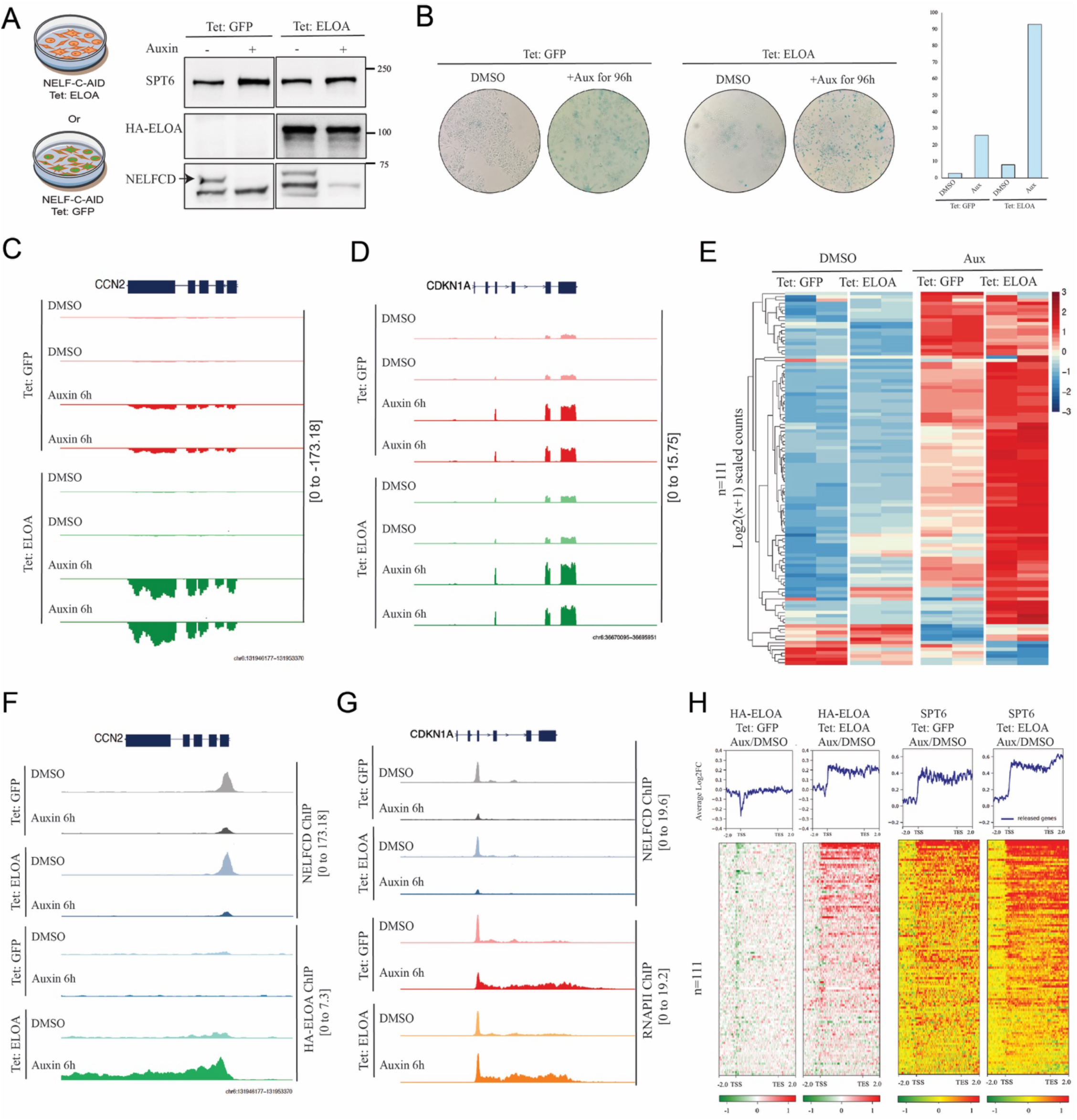
ELOA overexpression amplifies NELF-C depletion-induced gene activation. (A) Schematic view and western blots confirmation of generation of cell lines stably overexpressing N-terminal FLAG and HA-tagged ELOA on top of NELF-C-AID. Cells were treated with DMSO (-) or auxin (+) for 6h and Dox for 24h. (B) Staining for senescence-associated β-galactosidase activity in NELF-C-AID cells stably expressing GFP or ELOA and treated with DMSO or auxin for 96h. (C-D) Genome browser track visualizations of RNA-seq signal (two replicates are shown) at the senescence-associated genes CCN2 (C) and CDKN1A (D) in NELF-C-AID cells stably expressing GFP or ELOA and treated with DMSO or auxin for 6h. (E) Hierarchical clustering heatmap of gene expression levels (Log2-transformed, pseudocount-adjusted scaled RNA-seq read counts) in NELF-C-AID cells stably expressing GFP or ELOA and treated with DMSO or auxin for 6h, for cluster 1 genes (the set of genes at which RNAPII is strongly released upon auxin treatment in NELF-C-AID cells, as shown in Fig. 1F, n=111 genes). (F-G) Genome browser track visualizations of NELFC and HA-ELOA ChIP-seq signals at CCN2 (F), and NELFC and RNAPII at CDKN1A (G) in NELF-C-AID cells stably expressing GFP or ELOA and treated with DMSO or auxin for 6h. (H) Metaplots of Log2-transformed fold change (Log2FC, above) and heatmaps (below) of differential HA-ELOA and SPT6 ChIP-seq signal at the same set of genes as in (E), in auxin-treated relative to DMSO-treated NELF-C-AID cells stably expressing GFP or ELOA. TSS= transcription start site; TES=transcription end site.

### The primate-specific ELOA homolog ELOA3 plays a similar role upon NELF-C depletion

In addition to ELOA, our unbiased screen (Fig. 3 A-B) also identified the primate-specific ELOA homolog ELOA3 (TCEB3C) as a factor that might facilitate the growth arrest phenotype associated with NELF-C depletion. We validated this using clonal ELOA3 KO cell lines in the NELF-C-AID DLD-1 background, which we generated as described for ELOA KO using independent ELOA3-targeting sgRNA (Fig S6A). The results of our ChIP-seq experiments in ELOA-overexpressing cells suggested that the presence of NELF at promoters might preclude the association of ELOA with RNAPII or associated elongation factors, such as SPT6. However, we previously reported that ELOA3 interacts strongly with NELF, suggesting distinct interactions between NELF and either ELOA or ELOA3 (*42*). Therefore, to investigate the impact of NELF depletion on ELOA3 function, we stably overexpressed an N-terminal FLAG- and HA-tagged ELOA3 construct in NELF-C-AID cells (Fig. S6B). Next, we performed genome-wide sequencing experiments following Dox-induced overexpression of ELOA3. Similar to ELOA, overexpression of ELOA3 amplified NELF-C depletion-induced activation of CCN2, CDKN1A, and the rest of the genes in cluster 1 (Fig. S6D-F and Fig. S7A-C). Also similar to ELOA, overexpression of ELOA3 amplified the NELF-C depletion-induced occupancy of SPT6 throughout the body of CCN2, CDKN1A, and cluster 1 genes (Fig. S6G-I). However, unlike ELOA, overexpression of ELOA3 dramatically increased NELF-C occupancy at the CCN2 and CDKN1A gene promoters in untreated NELF-C-AID cells (Fig. S6G-H). This is consistent with our previous findings that ELOA3 and NELF-C interact strongly and colocalize at gene promoters (*42*). HA-ELOA3 also occupied the promoters of cluster 1 genes, and this occupancy was not lost upon NELF-C depletion (Fig. S6I, Fig. S7D). Instead, NELF-C depletion increased HA-ELOA3 occupancy within the gene body of cluster 1 genes (Fig. S6I, Fig. S7D), similar to the impact of NELF-C depletion on HA-ELOA (Fig. 4F-H, Fig. S5E). When comparing the number of genes that are upregulated upon NELF-C depletion combined with ELOA or ELOA3 overexpression, we see a significant overlap between ELOA and ELOA3 (Fig. S7E, Fig. S7F). Together, these results indicate that although ELOA3 differs from ELOA in its promoter-proximal interaction with NELF, this interaction is not required for chromatin occupancy, and ELOA3 can play a similar role to ELOA upon NELF-C depletion.

### ELOA loss suppresses NELF depletion-induced RNA processing defects

Given the identification of ELOA in our CRISPR screen and the impacts of ELOA loss observed via short-read RNA-seq, we sought to determine at single-molecule resolution whether loss of ELOA could rescue NELF-C depletion-induced changes in transcript isoform usage. To do so, we performed Iso-seq on matched sgControl versus sgELOA cell lines treated with auxin or DMSO for 6h. Visualizing fold changes in the representation of specific transcript classes within the global transcriptomes of these cells, we observed changes in isoform usage indicating RNA processing defects upon NELF loss; however, ELOA loss had no significant impact on these effects globally (Fig. 5A). Compared to untreated sgControl cells, acute NELF depletion led to down-regulation of incomplete splice-match (ISM) mono-exonic transcripts (sgCtrl: log2-fold change [log2FC] = -0.485, p = 0.018; sgELOA: log2FC = -0.134, p = 0.509) and upregulation of intron retention events in both the fusion category (sgCtrl: log2FC = 0.391, p = 0.023; sgELOA: log2FC = 0.215, p = 0.212) and the novel not in catalog category (sgCtrl: log2FC = 0.342, p = 0.017; sgELOA: log2FC = 0.231, p = 0.106), in both ELOA WT and ELOA KO backgrounds (Fig. 5A). Furthermore, correlating the fold-changes in isoform usage upon NELF degradation in ELOA WT cells against those upon NELF degradation in ELOA KO cells revealed only a muted, non-significant anti-correlation (r = -0.322, p = 0.116), suggesting that ELOA loss is not sufficient to reverse the global transcriptomic consequences of NELF loss.

**Figure 5.**
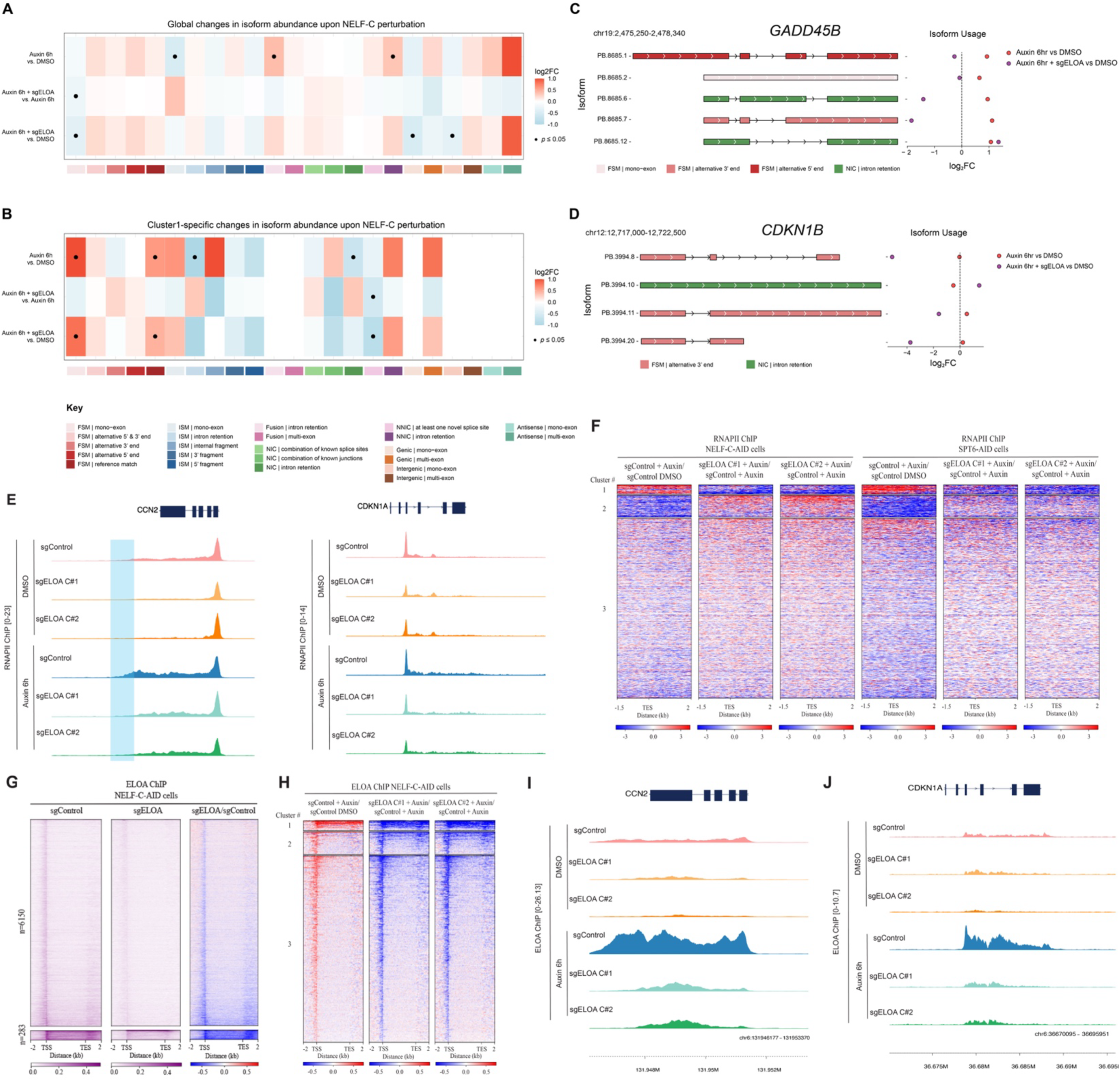
ELOA knockout partially rescues NELF-C depletion-induced transcriptional and co-transcriptional changes. (A-B) Global (A) and cluster 1-specific (B) changes in isoform class abundance upon NELF and/or ELOA depletion, quantified using DESeq2. Each panel shows the effects of NELF degradation in ELOA WT cells (Auxin 6h vs DMSO), ELOA KO on subsequent NELF degradation (Auxin 6h + sgELOA vs Auxin 6h), and NELF degradation in ELOA KO cells (Auxin 6h +sgELOA vs DMSO). We observe a significant anticorrelation between the effects of NELF degradation (in ELOA WT cells) and ELOA KO (on top of NELF degradation) for cluster 1 genes (Pearson’s r =-0.599, p = 0.007). (C-D) Track visualization of GADD45B (C) and CDKN1B (D) isoforms identified via Iso-seq, and changes in their usage upon NELF degradation in ELOA WT and ELOA KO cells. (E) Genome browser track visualization of RNAPII ChIP-seq signal at the senescence-associated genes *CCN2* and *CDKN1A* in sgControl and sgELOA (ELOA KO 1 and 2) NELF-C-AID cells treated with DMSO or auxin for 6h. (F) Genome-wide heatmaps of differential RNAPII ChIP-seq signal centered on transcription end sites (TES) in sgControl and sgELOA (ELOA KO 1 and 2) NELF-C-AID (left) and SPT6-AID (right) DLD-1 cells treated with DMSO or auxin for 6h. Clustering is based on differential RNAPII signal in NELF-C-AID cells; n=6433 genes, with n=286 genes in uppermost cluster. (G) Heatmaps of ELOA ChIP-seq signal in untreated sgControl and sgELOA NELF-C-AID DLD-1 cells, with heatmap of differential signal in sgELOA versus sgControl at right. 283 genes with the highest gene body and TES ELOA occupancy are shown at bottom. (H) Genome-wide heatmaps of differential ELOA ChIP-seq signal, for clusters defined in (F). (I-J) Genome browser track visualization of ELOA ChIP-seq signal at the senescence-associated genes *CCN2* (I) and *CDKN1A* (J) for the conditions in (E).

We next asked whether NELF depletion-induced RNA processing defects could be rescued by ELOA loss specifically for Cluster 1 genes (identified in Fig. 2E). Building on our previous analysis, we subset isoforms for Cluster 1 genes only, and repeated the same fold-change calculations as above. For these genes, we observed that acute NELF depletion led to down-regulation of intron retention events categorized as incomplete splice-match (ISM) (sgCtrl: log2FC = -1.574, p = 0.031; sgELOA: log2FC = -1.250, p = 0.087) or novel in catalog (NIC) (sgCtrl: log2FC = -0.658, p = 7e-4; sgELOA: log2FC = -0.332, p = 0.089), and up-regulation of full-splice match (FSM) mono-exonic transcripts (sgCtrl: log2FC = 1.145, p = 1e-4; sgELOA: log2FC = 0.827, p = 0.006), in both ELOA WT and ELOA KO backgrounds (Fig. 5B). Unlike the global phenotype, however, we found a striking and significant anticorrelation between the fold changes in sgELOA vs sgControl and those in auxin vs DMSO-treated cells (r = -0.599, p = 0.007). This suggests that for Cluster 1 genes, loss of ELOA rescues the transcriptome-scale defects in RNA processing (e.g. intron retention) induced by NELF loss. Specific examples of these changes induced by NELF degradation were visible on the genome browser (Fig. 5C-D).

Defects in RNA processing, including terminal exon definition and termination-coupled 3’ end cleavage and polyadenylation, may be reflected in transcription termination defects (*43*). Interestingly, when we examined NELF depletion-induced changes in RNAPII occupancy via ChIP-seq in WT or ELOA KO NELF-C-AID cells, we observed a marked and ELOA-dependent extension of RNAPII signal into the 3’ region of both *CCN2* and *CDKN1A* downstream of transcription end sites (TES) (Fig. 5E). Examining global RNAPII occupancy around TES in both NELF-C and SPT6 AID cells revealed a similar ELOA-dependent effect of NELF or SPT6 depletion for a group of 286 genes (Fig. 5F), which includes more than 90% of the genes with NELF depletion-induced RNAPII release (n=111, identified in Fig. 2E) (Supplement). When we analyzed ELOA occupancy via ChIP-seq in untreated NELF-C-AID cells, we identified a group of 283 genes that have high ELOA levels within the gene body and around the TES prior to elongation factor depletion (Fig. 5G). Remarkably, more than 84% of the genes with NELF depletion-induced RNAPII release (n=111, identified in Fig. 2E) are part of this group (Supplement). Finally, we examined the impact of NELF depletion on ELOA occupancy via ChIP-seq in WT or ELOA KO NELF-C-AID cells, observing elevated levels of ELOA occupancy at the 286 genes with ELOA-dependent, NELF depletion-induced increases in RNAPII occupancy around TES (Fig. 5H, gene set identified in Fig. 5F). Of note, ELOA occupancy was strongly increased by NELF depletion at the senescence-associated genes *CCN2* and *CDKN1A* (Fig. 5I-J). Together, these data indicate that 1) the genes where RNAPII is released upon NELF depletion are ELOA-occupied genes, and that 2) NELF-depletion induced 3’ RNAPII occupancy past TES is driven by dramatic increases in ELOA levels at these ELOA-occupied genes.

### Acute loss of ELOA results in a 3’ elongation defect that affects cellular senescence-associated cluster 1 genes

To further investigate the function of ELOA in the growth-arrest phenotype observed upon NELF depletion, we generated a degron line to enable acute depletion of ELOA using the dTAG system in the DLD-1 background (ELOA-dTAG cells). Rapid depletion of ELOA from ELOA-dTAG cells was achieved within 5 hours following addition of dTAG-13 to the culture medium (Fig 6A). We performed ChIP-seq to determine the impact of acute ELOA depletion on NELFCD chromatin occupancy. Interestingly, upon loss of ELOA from gene promoters, NELF occupancy at gene promoters also decreased genome-wide (Fig. 6B). When we clustered genes based on ELOA occupancy levels within gene bodies, we identified a group of genes (n=184) with particularly high gene body ELOA occupancy (Fig. 6C). There is significant overlap between this group of genes and cluster 1 genes, at which RNAPII is released upon NELFCD depletion (Fig. 2E, Fig. 6D). To analyze impacts of ELOA loss on transcription by RNAPII, we also performed ChIP-seq for RNAPII and its elongation-associated CTD modification Ser2P. For cluster A genes, we observed increased occupancy of elongating RNAPII within gene bodies that was accompanied by a striking loss of occupancy at the 3’ end of gene regions (past transcription end sites) upon acute ELOA depletion (Fig. 6E), consistent with defects in RNAPII processivity. We also observed this 3’ elongation defect at some senescence-associated genes such *CCN2* (Fig. 6F) and at some immediate early genes such as *EGR1* (Fig. 6G), which suggests a potential role for ELOA in regulating stress response. Interestingly, genes with high gene body ELOA occupancy = are all short genes <30kb (Fig. 6H), further validating the potential role of ELOA in stress response. Overall, these data are consistent with those in Figure 5 and lend insight into the function of ELOA beyond the artificial condition of NELF depletion.

**Figure 6.**
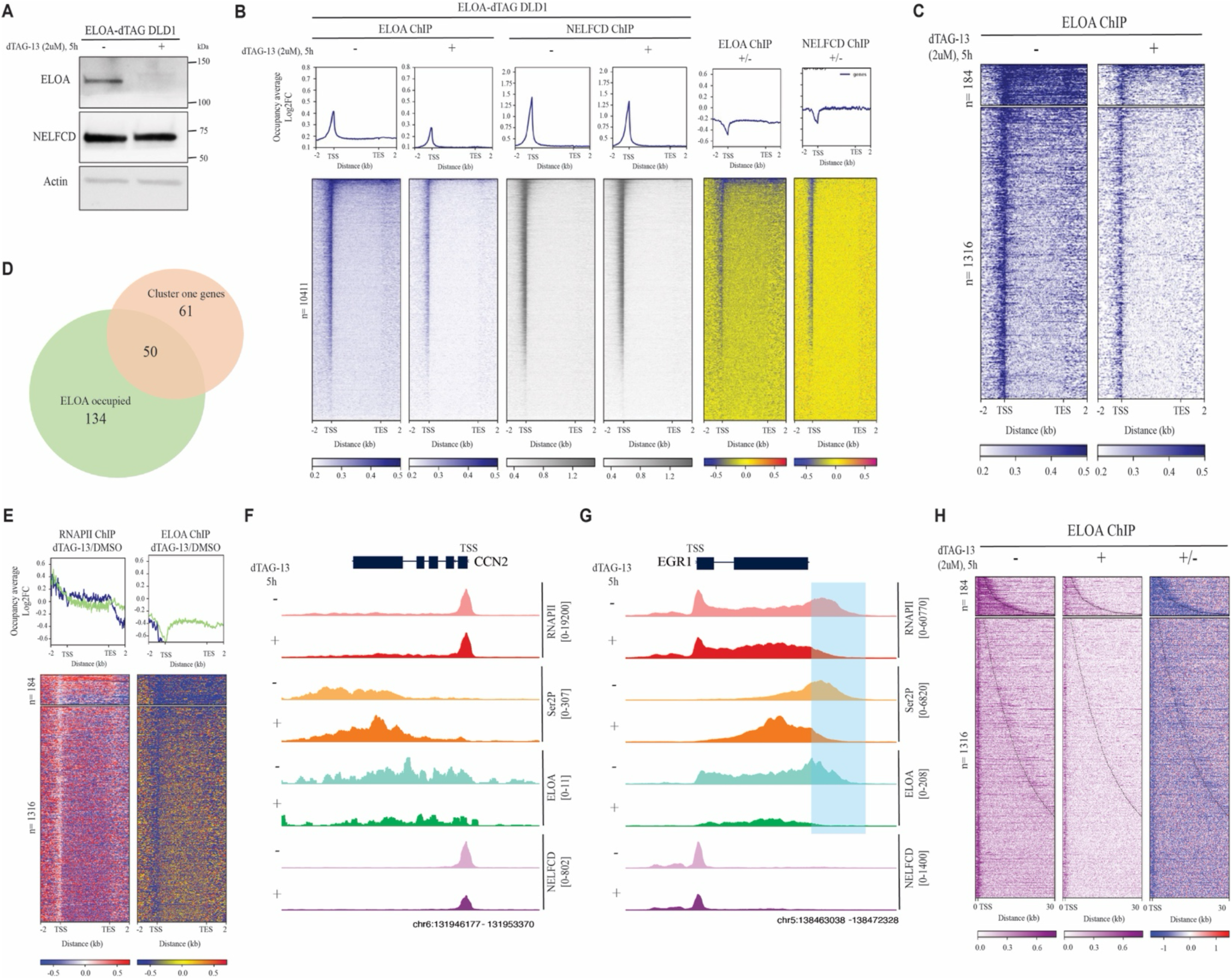
Acute loss of ELOA via dTAG results in 3’ elongation defects. (A) Western blot analysis of ELOA depletion upon dTAG-13 treatment (2uM, 5h) in the ELOA-dTAG DLD-1 cell line. Blots were probed for ELOA and NELFC with actin as loading control. (B) Genome-wide metaplots of Log2-transformed fold change (Log2FC, above) and heatmaps (below) of differential ELOA and NELFCD ChIP-seq signal in ELOA-dTAG DLD-1 cells upon treatment with DMSO or dTAG-13 (2uM, 5h). (C) Heatmaps of differential ELOA ChIP-seq signal in ELOA-dTAG DLD-1 cells (treated as in B) at the top 1500 genes with highest gene body ELOA occupancy, from which a small cluster of genes (n=184) that have particularly high gene body ELOA occupancy was identified. (D) Venn diagram illustrating the overlap between cluster 1 genes (n=111, genes released upon NELFCD depletion, see Fig. 2E) and genes that have high ELOA occupancy within the gene body (n=184). (E) Genome-wide metaplots of Log2-transformed fold change (Log2FC, above) and heatmaps (below) of differential ChIP-seq signal (RNAPII, Ser2P, ELOA and NELFCD) at genes with high ELOA occupancy (shown in C). (F-G) Genome browser track examples of ChIP-seq signal (RNAPII, Ser2P, ELOA and NELFCD) at the example genes *CCN2* (F) and *EGR1* (G). (H) Heatmaps of differential ELOA ChIP-seq signal in ELOA-dTAG DLD-1 cells (treated as in B) at the top 184 genes with highest gene body ELOA occupancy, are shown from TSS to 30kb distance from the TSS.

### ELOA loss confers a growth advantage to aging human primary dermal fibroblasts

Our data indicate that auxin-induced depletion of NELF or SPT6 results in a growth arrest phenotype that involves expression of senescence-associated genes and is mediated by ELOA. To investigate potential function of ELOA in a physiological condition, we chose to use human dermal primary fibroblasts as a model of cellular senescence in natural aging. We initially generated ELOA knockout in these cells at passage 1 via two independent sgRNA (Fig. 7A); because ELOA depletion was more efficient in cells expressing sgRNA#2, we then examined the impact of ELOA loss over an extended period in these cells in comparison to cells expressing a non-targeting sgRNA (sgControl). By passage 16, p21 protein levels were reduced in ELOA KO relative to control fibroblasts, and capacity for cell growth (assessed as total number of crystal violet-stained cells at 11 days versus 1 day post-seeding) was significantly increased (Fig. 7B-F). RNA-seq analysis confirmed that expression of CCN2 and CDKN1A (p21) was reduced at the mRNA level in ELOA KO relative to control fibroblasts by passage 16 (Fig. 7G). We also analyzed expression of genes within the SenMayo gene set (*30*), which was previously reported to identify senescent cells across human tissues. We found that more than half of these genes were downregulated in ELOA KO compared to control fibroblasts at passage 16 (Fig. 7H), which could potentially contribute to the growth advantage conferred by loss of ELOA. To corroborate our RNA-seq data, we performed CUT&RUN for the elongation-associated RNAPII CTD modification Ser2P in early versus late passages (passage 7 versus passage 17) of both sgControl and sgELOA fibroblasts. Consistent with RNA-seq, Ser2P signal increased in an ELOA-dependent manner over the *CCN2* gene body in late versus early passage fibroblasts (Fig. 7I). This pattern was also observed for cluster 1 genes with Ser2P signal centered around TES (Fig. 7J). These data support a potential role for ELOA in the regulation of senescence and the progression of aging in human cells.

**Figure 7.**
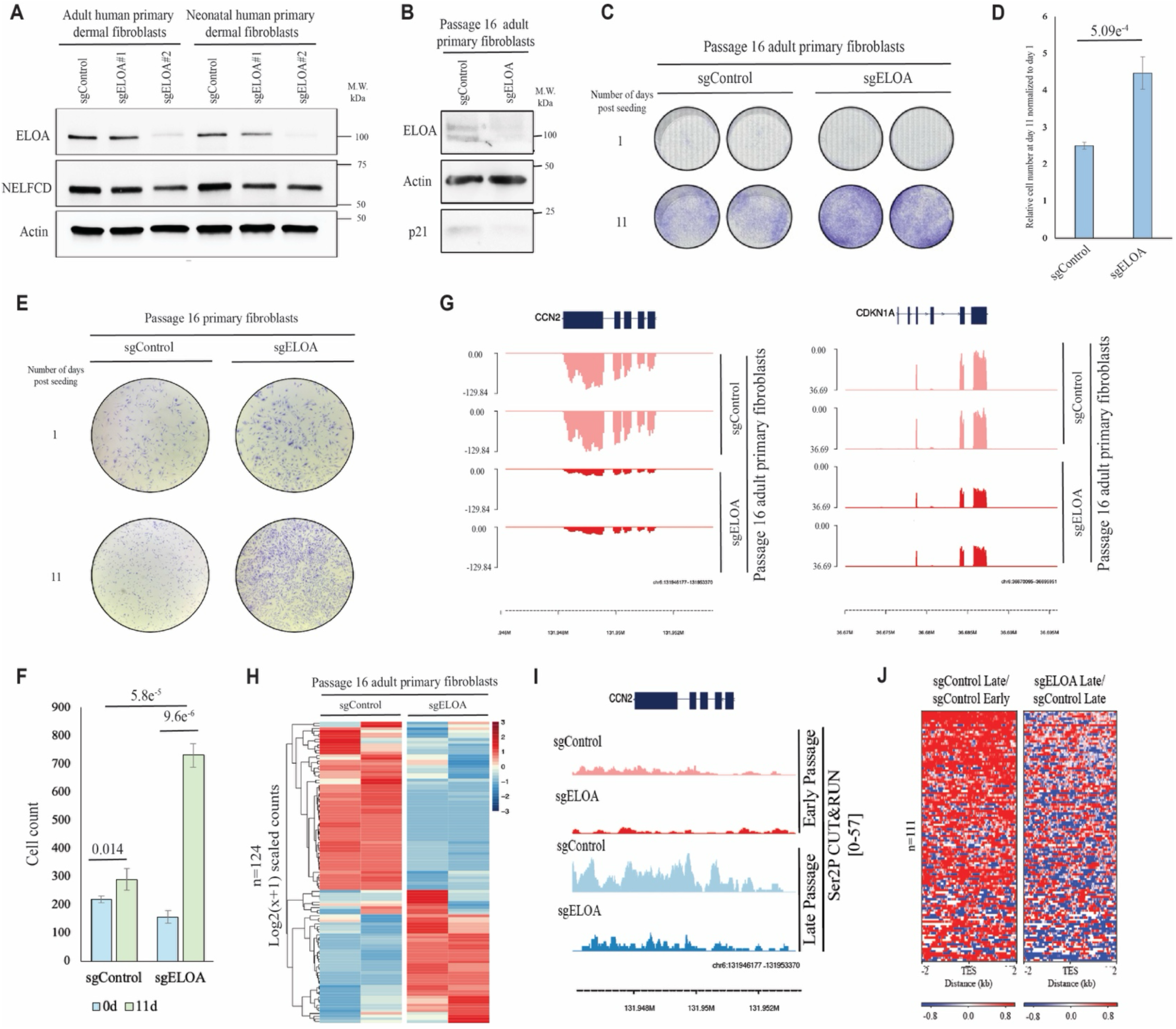
ELOA knockout confers a growth advantage in human primary dermal fibroblasts. (A) Western blot analysis of ELOA depletion upon pool ELOA KO using the same independent sgRNAs as in Fig. 2 (sgRNA#1 and sgRNA#2), with a nontargeting sgRNA as control (sgControl), in both adult and neonatal human primary dermal fibroblasts. Blots are probed for ELOA and NELFCD with actin as loading control. (B) Western blot analysis of ELOA depletion in ELOA KO adult human primary dermal fibroblasts (primary fibroblasts) and abated p21 expression at passage 16. (C-E) Accelerated growth in ELOA KO (sgELOA) compared to sgControl primary fibroblasts (C), with manual quantification in (D), magnified view in (E), and automated quantification via ImageJ in (F). All p-values in (D) and (F) are calculated using student t-test. (G) Genome browser track visualization of RNA-seq signal at the senescence-associated genes CCN2 and CDKN1A in wild-type (sgControl) and ELOA KO (sgELOA) primary fibroblasts at passage 16. (H) Differential expression heatmap for SenMayo signature genes (n=124) in ELOA WT vs KO primary fibroblasts at passage 16. The complete list of these genes can be found in Supplemental files. (I) Genome browser track visualization of Ser2P CUT&RUN signal at the *CCN2* locus in wild-type (sgControl) and ELOA KO (sgELOA) primary fibroblasts at passage 7 (early) and passage 17 (late). (J) Heatmaps of differential Ser2P CUT&RUN signal in sgControl primary fibroblasts at passage 17 (late) relative to passage 7 (early), and in sgELOA relative to sgControl primary fibroblasts at passage 17 (late), for cluster 1 genes (genes released upon NELFCD depletion, see Fig. 2E). TES=transcription end site. n=111 genes.

## Discussion

Accurately mapping the molecular connections between aspects of RNA synthesis and processing (*i.e.* transcription initiation, elongation, termination, and splicing) promises novel insights into how genes are regulated. Here, we have presented a single-molecule transcriptomic ‘perturbation atlas’ spanning four genetically engineered degron cell lines targeting the essential elongation factors PAF1, SPT5, SPT6, and NELF. This atlas – generated using the PacBio Iso-Seq method and quantified using MPAQT, a state-of-the-art algorithm that integrates short- and long-read RNA-seq data – reveals new dimensions of transcriptome space and offers an invaluable glimpse into how RNA processing is directly regulated by core elongation factors. In addition to finding that specific isoforms, genes, and pathways are distinctly impacted upon degradation of these different factors, we have also identified an intriguing overlap within NELF and SPT6 datasets, showing upregulation of isoforms and genes involved in cellular senescence. Beyond their use here, these datasets will be particularly valuable to computational biologists requiring ‘perturbation’ training data for the next generation of AI models for predicting RNA splicing.

In vitro biochemical studies have long suggested that NELF is a factor regulating the promoter-proximal pausing of transcriptionally engaged RNAPII, a checkpoint that also allows for rapid unleashing of transcriptional programs in response to cell intrinsic and environmental cues (*15, 44, 45*). Our group provided new insight into NELF function by demonstrating that acute depletion of NELF from mammalian cells does not result in a genome-wide release of RNAPII into the gene body, as might be expected of a “negative” elongation factor (*15*). Instead, NELF depletion results in genome-wide stalling of RNAPII at a downstream region associated with the +1 nucleosomal dyad, in a transition that occurs independently of the positive transcription elongation factor PTEFb (*15*). Here, we have disclosed an exception to this genome-wide pattern of second-site RNAPII pausing: for a small set of genes (less than 2% of expressed genes, referred to as cluster 1 genes, Fig. 2D-E), we observed strong release of RNAPII into the gene body upon NELF depletion (Fig. 8A-B). These include the genes *CDKN1A*, *CCN2*, and *ID2*, which are all strongly associated with or implicated in the initiation of cellular senescence. CCN2 is of particular interest as a senescence-associated secreted factor that has previously been shown to induce senescence in a paracrine manner (*35*). We have also shown that NELF loss results in β-galactosidase activity, a bona-fide senescence-associated marker. These findings might appear to indicate similarity between the NELF depletion-induced growth arrest phenotype and cellular senescence, which is typically thought to involve irreversible growth arrest. However, the NELF depletion-induced growth arrest phenotype is reversible upon auxin washout, indicating potential similarity to cellular quiescence (a state of reversible growth arrest). Whether quiescence and senescence are qualitatively distinct or whether they exist on some continuum of quiescence “depth” is a matter of debate (*46*) and recent results support the conclusion that quiescence and senescence are molecularly indistinguishable at any single point in time (*47*).

**Figure 8.**
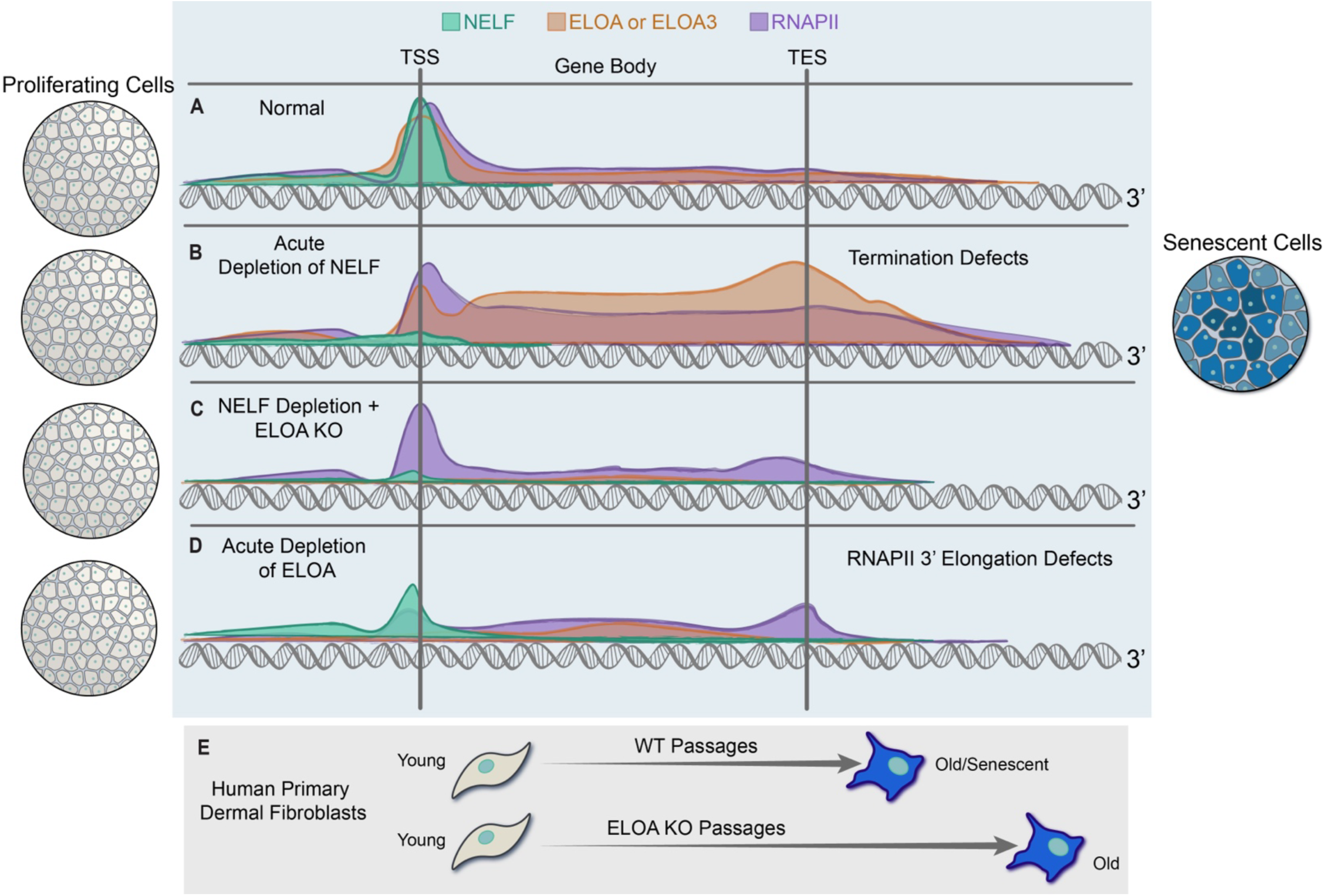
Functional NELF and ELOA interaction with RNAPII in cellular senescence/quiescence. Our study reveals the functional interaction of NELFC and ELOA in elongation control at senescence-associated genes. Acute depletion of the critical transcription factor NELF results in a reversible growth arrest phenotype. This is accompanied by the release of RNAPII at a small number of genes (cluster 1) including genes associated with cellular senescence/quiescence, such as CDKN1A (p21) and CCN2. (A) Under normal conditions, NELF and ELOA associate with RNAPII at the promoters of cluster 1 genes. (B) Acute depletion of NELFC dramatically increases ELOA levels at cluster 1 genes, driving ELOA-mediated transcription that extends past transcription end sites; the resulting transcripts reflect RNA processing and termination defects. (C) Loss of ELOA rescues the NELF depletion-induced transcript defects and growth arrest phenotype. (D) When NELF levels are normal, acute ELOA depletion results in marked defects in transcriptional elongation towards the 3’end of otherwise ELOA-occupied genes. (E) Consistent with ELOA function in regulating cellular senescence, human dermal primary fibroblasts lacking ELOA show an enhanced capacity for cell growth at higher passage numbers.

Our findings also raise the possibility of a direct role for NELF in the regulation of cellular senescence and aging. This would be consistent with a previous report indicating a role for NELF dosage in the regulation of longevity in Drosophila via control of transcription at heat shock protein genes (*48*). Further studies are needed to determine how different components of NELF dosage such as expression, stability, and differential interaction with RNAPII and/or other elongation factors might impact this potential role. Via genome-wide CRISPR screens, we have identified a role for the elongation factor ELOA as a critical mediator of growth arrest induced by loss of NELF or SPT6. ELOA KO rescued the effects of NELF depletion, and ELOA overexpression amplified these effects, including increased expression of senescence-associated genes. While ELOA can enable gene expression in the absence of NELF, our further analyses indicate that this comes at the cost of defects in transcription termination (Fig. 8C), which is coupled to 3’ transcript processing. We have identified global shifts in transcript isoform usage indicating RNA processing defects in NELF-depleted cells. It is important to note that because the Iso-Seq protocol is designed to profile steady-state and poly(A)mRNA, our findings likely underestimate the consequences of NELF depletion and ELOA reintroduction, given the average half-life of human mRNAs of ∼10 h (*49*). Moreover, our dataset is not likely to capture nuclear RNA processing events prior to cleavage and polyadenylation. Age-related changes in transcript isoforms resulting from alternative splicing have previously been reported in different cell types and animal models (*50–52*). Intron retention in particular has been characterized as a key fingerprint of aging in C. elegans, mice, and Drosophila (*53–55*). Despite a number of reports implicating splicing defects in aging-related phenomena, mechanisms have remained elusive. Investigating the crosstalk between NELF, ELOA, and other factors in the regulation of pre-mRNA splicing may be critical to understanding cellular senescence and aging.

Pre-mRNA splicing is not only functionally and spatially but also kinetically coupled to transcription. Even post-transcriptional splicing is influenced by RNAPII processivity (a component of elongation rate) due to the impact of elongation rate on the conformation of the nascent transcript, which alters its availability to different spliceosomal components and other RNA binding proteins including those involved in termination-coupled 3’ transcript processing (*56*). We have shown that acute ELOA depletion induces a 3’ elongation defect consistent with loss of processivity or premature termination at genes where ELOA accompanies elongating RNAPII into the gene body under normal conditions (Fig. 8D). This is consistent with a physiologic role for ELOA in maintaining RNAPII processivity, or potentially in termination-coupled 3’ transcript processing, although this latter role may be played indirectly via impacts on NELF.

Given the long-debated role of ELOA as an elongation factor (*1, 5, 10, 11, 13, 57*), the gene expression data we have presented for ELOA KO, ELOA overexpression, and acute ELOA depletion shed light on its capacity to function in transcriptional elongation control while also suggesting a likely novel function for ELOA in the control of processes involved in cellular senescence (or quiescence). Remarkably, over 90% of the genes at which RNAPII is released upon NELFCD depletion (cluster 1 genes), including genes implicated in cellular senescence, are genes that are occupied by ELOA under normal conditions. While our data indicate that ELOA can support RNAPII elongation to sub-optimally “supplement” for loss of NELF, we have also observed that loss of ELOA leads to loss of NELF from promoters. This appears to occur independently of the level of RNAPII promoter occupancy, indicating that ELOA may be able to regulate NELF levels in a highly localized (gene locus-specific) manner. Future studies may determine how ELOA and/or NELF dosage may change during aging, and what factors facilitate gene specificity for ELOA. Employing human primary dermal fibroblasts as a cell-culture based model of natural human aging, we have observed a strong growth advantage for ELOA KO relative to wild-type at late passages (Fig. 8E). Intriguingly, we have also demonstrated that despite differing from ELOA in its interaction with NELF (*42*), the primate-specific ELOA homolog ELOA3 is able to play a similar role upon NELF depletion. However, further work is needed to delineate the mechanistic interplay between RNAPII, NELF, and ELOA (or ELOA3). Future biochemical studies will determine how ELOA, ELOA3 and different NELF subunits interact with each other and with other RNAPII-associated factors such as SPT6, whether and under what conditions they compete for binding to RNAPII, and how these interactions may impact cell fate under stress conditions or upon aging.

## Acknowledgements

We thank Marc A. Morgan for insights and resources and Brianna M. Monroe for graphical visualization.

## Author Contributions

Conceptualization: AS, SP

Methodology: SP, YA, VR, MI

Software: MI, SP, SW, DO, VR

Formal analysis: MI, SP, DO, SW, VR

Investigation: SP, YH, JZ, WT, BH, SW

Resources: AS, VR, YA

Data curation: MI, DO, SP, SW

Writing – original draft: SP, SG, SW

Writing – review and editing: SG, SP, VR, MI

Visualization: MI, DO, SP, SW

Supervision: AS

Funding Acquisition: AS, VR

## Supplementary Materials

## Materials and Methods

### Cell Culture

HEK293T (CRL-3216) and DLD-1 (CCL-221) cells were acquired from ATCC. Information on NELF-C-AID and NELF-E-AID cells can be found in a previous publication from our group (*15*). All cells were cultured in Dulbecco’s modified Eagle’s medium (DMEM) from Thermo Fisher (cat# 11965-118) supplemented with 15% fetal bovine serum (FBS) from Millipore Sigma (cat# F2442), 1x Glutamax from Thermo Fisher (cat# 35050-079) and 100 units/ml penicillin-streptomycin from Thermo Fisher (cat# 15140-163). To select lentivirus-infected cells, puromycin (1.5 μg/ml) and hygromycin (250 μg/ml) were used. All cells were tested for mycoplasma contamination using a MycoStrip detection kit (Invivogen, rep-mys-100) and Myco PCR with primers listed in Supplemental Table 2.

### Human Primary Fibroblasts

Adult (PCS-201-012) and Neonatal (PCS-201-010) Human primary dermal fibroblasts were bought from ATCC. These cells were cultured in Fibroblast Basal Medium (ATCC, PCS-201-030) containing components (L-glutamine: 7.5 mM, rh FGF basic: 5 ng/mL, rh Insulin: 5 µg/mL, Hydrocortisone: 1 µg/mL, Ascorbic acid: 50 µg/mL, Fetal bovine serum: 2%) of a low serum kit (ATCC, PCS-201-041). These cells were tested for mycoplasma regularly using the primers listed in Supplemental Table S1. Pool ELOA knockout was generated in these cells using two independent sgRNAs (listed in Supplemental Table S1) cloned into lentiCRISPR v2 puro. To select for ELOA KO in these cells, puromycin (1.5 μg/ml) was used. Upon selection, these cells were passaged regularly until passage 16, where protein and RNA were obtained from these cells for downstream western blot and RNA-seq. For growth assay in these cells, equal number of cells (25K) from the passage 16 of both WT and ELOA KO adult human primary fibroblasts were seeded (two 24-well plate for each) in triplicate. A day after one plate was fixed and another plate was fixed 11 days after the initial seeding. Growth (Quantification of the crustal violet) at day 11 was normalized to day 1. Cells were also counted using ImageJ by setting a color threshold, selecting for hue in the purple range, and then analyzing particles based on size.

### dTAG lines generation

sgRNAs (targeting stop codons of ELOA or NELF) for generation of knock in lines (ELOA-dTAG in DLD-1 and SYO-1, NELFC-dTAG in SYO-1 and NELFC- and ELOA double dTAGs in SYO-1) are listed in Supplemental Table S1. These sgRNAs were cloned into PX458 (Addgene, Plasmid #48138). The donor plasmid containing the HA-FKBP12^F36V^, the homology arms and resistancy genes (Blasticidin for ELOA and Neomycin for NELF-C) was designed and ordered from Twist Biosciences (sequence verified) in pTwist plasmid backbone (pTwist Amp High Copy). Transfection was done using Lipofectamine™ 3000 Transfection Reagent (Thermo Fisher Scientific, L3000001). Two days post transfection, we started to select cells with Blasticidin for ELOA-dTAG and Neomycin for NELFC-dTAG. Upon successful selection for 5 days, cells were plated at a density of 500 cells per 10cm dish for clonal generation of homozygous ELOA or NELFC -dTAG. Once NELF-C-dTAG was generated, we started to make an ELOA dTAG in that cell line to generate the double dTAG cell line (NELFC- and ELOA double dTAGs in SYO-1).

### Lentivirus production

HEK293T cells were seeded in 10cm or 15cm dishes at a density of 100,000 cells per cm² one day before transfection. One day later, the culture medium was replaced with fresh medium. Over the next 2 days, the medium containing lentiviral particles was collected daily and stored at 4°C. The lentiviral medium was then filtered through a 0.45 μm filter and concentrated using a PEG (polyethylene glycol) precipitation solution. The mixture was then incubated overnight at 4°C on a rocking platform to allow lentiviral particles to precipitate. The following day, centrifugation separated the viral particles (pellet) from the solution (supernatant). The supernatant was discarded, and the pellet containing the lentiviral particles was resuspended in sterile PBS.

### Plasmids

sgRNA expression constructs were made by inserting sgRNA sequences directly into lentiCRISPR v2 plasmids using a Quick Ligation Kit (New England Biolabs). For Doxycycline-inducible expression constructs DNA G-block fragments containing the desired genes were first assembled using NEBuilder HiFi DNA Assembly. These assembled fragments were then inserted into pCW57-MCS1-P2A-MCS2 plasmids that had been cut with specific restriction enzymes (NheI-HF and SalI-HF). lentiCRISPR v2 (Addgene, #52961) was a gift from Feng Zhang. pCW57-MCS1-P2A-MCS2 (Hygro) (Addgene, #80922) was a gift from Adam Karpf. pMD2.G (Addgene, #12259) and psPAX2 (Addgene 12260) were gifts from Didier Trono. Human Brunello CRISPR knockout pooled library (Addgene, #73179) was a gift from David Root and John Doench (*59*).

### CRISPR screening

The CRISPR screen methodology used here was described in our recent paper (*42*). The amplification of the Brunello CRISPR library in LentiCRISPR V2 (Plasmid #52961) was done using Endura electrocompetent cells (Lucigen, 60242-1) according to the manufacturer’s recommendation. NELF-C-AID or parental DLD-1 cells (1×10^7^ cells per 15 cm dish) were infected with a multiplicity of infection of approximately 0.5. Cells were infected in 3 separate batches as replicates, each batch contained two large dishes (15 cm) with 2×107 cells per dish. After one day, the medium was removed and replaced with fresh growth medium. Two days post-infection, cells were re-plated at a slightly lower density (1.5×10^7^ cells per 15cm dish) with fresh growth medium containing 10ug/ml puromycin. To select for cells that had successfully incorporated Brunello library, cells were treated with puromycin for 6 days and then plated at a density of 1×10^7^ million cells per 15 cm dish, with six 15 cm dishes (three replicates of two plates each) each for parental and NELF-C-AID DLD-1 cells. The following day auxin at a concentration of 1 micromolar (uM) was added to each plate. The growth medium was replaced with fresh auxin-containing medium every day. To maintain a similar cell number throughout the experiment, parental DLD-1 cells were passaged every two days at a density of 1×10^7 cells per dish. After 10 days of selection with auxin treatment, cells were collected by trypsinization and counted. The parental DLD-1 samples yielded approximately 5×10^7^ cells per replicate and the NELF-C-AID DLD-1 samples yielded approximately 4×10^7^ cells per replicate. Genomic DNA was extracted from each replicate using the Quick-DNA Midiprep Plus Kit (Zymo, D4075). For sequencing library preparation, sgRNA sequences were amplified from genomic DNA (up to 10ug per 100ul reaction) using Q5 Hot Start High-Fidelity 2X Master Mix according to the manufacturers protocol (New England Biolabs, M0494L) and primers listed in Supplemental Table S1. The following PCR cycling conditions were used: initial denaturation at 98°C (30 seconds); 28x amplification cycles comprised of denaturation at 98°C (5 seconds), primer annealing at 63°C (20 seconds), and extension at 72°C (30 seconds); and final extension at 72°C (2 minutes). The resulting libraries did not require gel purification. Libraries were then pooled and sequenced on an Illumina NovaSeq 6000 instrument as single end 50b reads. For analysis, MAGeCK (*60*) count and test commands were run using default parameters with non-targeting sgRNAs used for normalization. An exactly similar strategy was followed for the CRISPR screening experiment in SPT6-AID DLD1 cells.

### ChIP-seq

Buffer formulations for ChIP experiments are described somewhere else (*42, 61*). Briefly, about 5×10^7^ cells were crosslinked by 1% formaldehyde at a density of 1×10^7^ (Millipore Sigma, F8775) for 15 minutes at room temperature. Then glycine was used to quench formaldehyde to 0.125M concentration for 5 minutes at room temperature. Next, the cells were washed with PBS twice each time pelleted at 2,000xG, snap frozen in liquid nitrogen and stored at –80 for further processing. For ChIP, lysis buffer 1 was used to lyse the cells at a density of 1×10^7^ cells per ml and incubation for 10 minutes on ice. After centrifugation at 2,000xG, the resulting nuclei were resuspended in lysis buffer 2 at a density of 1×10^7^ cells per ml and incubated on ice for 10 minutes. Next, once the nuclei were pelleted at 2,000xG, they were resuspended in lysis buffer 3 at a concentration of 5×10^7^ cells per ml. Subsequently, the nuclei were transferred to 1ml milliTUBEs with AFA fibers (Covaris, 520130) and sonicated in a Covaris E220 focused ultrasonicator (duty factor: 10%, peak intensity pulse: 140, cycles per burst: 200, time: 500 seconds). After pelleting the insoluble material at 20,000xG, using a DC protein assay kit (Bio-Rad, 5000112) protein concentration in the supernatant fraction was quantified relative to a standard curve of bovine serum albumin dissolved in Buffer 3. Protein concentration across the samples was normalized to 1-2 mg/ml and then Triton X-100 was added to a final concentration of 1%. Antibodies were added (see Supplemental Table S2) and samples were incubated overnight at 4°C on an end-over-end rotator. The following day, 30ul of washed Protein G Dynabead slurry (Thermo Fisher Scientific, 10003D) (pre-washed with Buffer 3 plus 1% Triton X-100) was added to each sample and the samples were incubated at 4°C on an end-over-end rotator for another 2 hours. Then, samples were then placed in a DynaMag magnet stand (Thermo Fisher Scientific, 12321D) and washed with 1ml of Wash Buffer (RIPA) 4x 5 min and with TE+NaCl 2x 5 min. Elution was performed using 200ul of Elution Buffer and incubation at 65°C for 2 hours. After removing the Dynabeads from the eluted immunoprecipitated chromatin on a DynaMag stand, proteinase K was added at a concentration of 0.5ug/ml prior to incubation overnight at 65°C. Next, DNA was purified using a ChIP DNA Clean and Concentrator kit (Zymo, D5205). Libraries were made using the KAPA HTP Biosystems Library Prep kit (Roche, KK8234) using Perkin Elmer Unique Dual Index Barcodes for NovaSeq 6000. Sequencing was done on an Illumina NovaSeq 6000 instrument as paired-end 50b reads. Reads were aligned to the hg38 human genome assembly using Bowtie version 1.1.2 (*62*). Peaks were called using MACS2 (*63*) and occupancy plots were generated using DeepTools (*64*).

Briefly, RNAPII release was assessed by calculating the pause-release ratio (PRR), which is the ratio of RNAPII occupancy level over the close gene body to that of the promoter region. Full gene body coverage ratio (FGBC) is calculated as RNAPII occupancy over the length of a full gene body, which is an average per base occupancy within the gene body region, calculated as TSS+300 to TES. All ChIP experiments were done once unless otherwise stated in the manuscript.

### CUT&RUN in fibroblasts

A day before the CUT&RUN experiments, 5×10^4^ sgControl or sgELOA from both early (passage 7) or late (passage 17) passages of human primary dermal fibroblasts were seeded in a 6cm dish. The day after cells were washed twice with cold 1xPBS at 4C and then the CUT&RUN Assay Kit from Cell Signaling Technology (#86652) was used to perform the CUT&RUN experiment following the protocol they described in detail (https://www.cellsignal.com/products/cut-run-kits-reagents/cut-run-assay-kit/86652?srsltid=AfmBOopLpRYTXFkhlIynj-LBmOsO3Mbvt1qHrEYxRzA8c7qR289veDhB). Sequencing was done on an Illumina NovaSeq 6000 instrument.

### GO terms analysis

Functional analysis was performed with g:Profiler (*65*). Multiple testing correction performed by g:SCS algorithm. It corresponds to an experiment-wide threshold of a=0.05, i.e. at least 95% of matches above threshold are statistically significant.

### RNA-seq

RNA was extracted from cells using the RNeasy Mini Kit (Qiagen, 74106) using an on column DNaseI digestion step (Qiagen 79256). RNA was subjected to polyadenylated RNA enrichment using the NEBNext Poly(A) mRNA Magnetic Isolation Module Kit (New England Biolabs, E7490L), and libraries were synthesized using the NEBNext Ultra II Directional RNA Library Prep Kit for Illumina (New England Biolabs, E7760L). RNA-seq libraries were sequenced on an Illumina NovaSeq 6000 instrument using single-end 50 cycle reads. Reads were aligned to the hg38 human genome assembly. Sequence quality was assessed using FastQC v 0.11.2, and quality trimming was done using Trimmomatic (*66*) with parameters TRAILING:30 MINLEN:20. RNA-seq reads were aligned to the hg38 genome using STAR v.2.5.2 (*67*) and only uniquely mapped reads with a two-mismatch threshold were considered for downstream analysis. RNA-seq reads were quantified to the gene level with HTSeq version 0.6.0 (*68*). Batch correction was done using RUVseq (*69*) and differential gene expression analysis was performed using DESeq2 (*70*). All RNA-seq experiments were done in two or three biological repeats.

### Long-read sequencing

For prepping RNA samples for long read, standard Pacbio Iso-seq protocol was followed (https://www.pacb.com/wp-content/uploads/Procedure-checklist-Preparing-Iso-Seq-libraries-using-SMRTbell-prep-kit-3.0.pdf. Brielfy, 300 ng purified RNA per sample was used for Pacbio Iso-seq library preparation, which includes cDNA synthesis, cDNA amplification, repair and A-tailing, SMRTbell adapter ligation, and exonuclease treatment. To assess library quality, samples were run on Agilent Bioanalyzer DNA chips or Agilent Femto Pulse. Libraries were sequenced on either Sequel II or Revio flow cells. Sequencing reads were then processed with software from Pacific Biosciences. Circular consensus sequences (CCS) were generated from raw subreads from the Sequel II system using ccs or directly obtained from the Revio system. CCS were demultiplexed using lima. Reverse transcription primer sequence flanking the mRNA sequences was removed using lima to generate full-length reads. Concatemers and polyA tails were removed using isoseq3 to generate full-length non-concatemer reads. Full-length non-concatemer reads were grouped into read clusters using hierarchical clustering in isoseq cluster2. High quality reads were aligned to the reference genome using pbmm2, and then converted into collapsed aligned reads using isoseq collapse. Statistical analysis on collapsed reads was performed via SQANTI3 package. Further statistical analyses were conducted in bulk and in selected clusters of genes indicated by bulk RNA-seq.

### Growth assay

Cells were seeded at a density of 50,000 cells per well in quadruplicate in 24 well plates in 1ml of growth medium. The following day, the medium was replaced with Auxin (500uM) or DMSO containing medium. Fresh medium containing Auxin or DMSO was added to the cells every day for three days. After 4 days, medium was aspirated, and cells were fixed in 4% formaldehyde (Millipore Sigma, F8775) in PBS (Millipore Sigma, P5358-10PAK) for 20 minutes at room temperature. Next, formaldehyde solution was decanted, and the plates were washed in tap water (Department of Water Management, Chicago, IL) and then air-dried overnight. Next, 1ml crystal violet solution (Millipore Sigma, HT90132-1L) was used to stain the cells for 1 hour at room temperature, after which the staining solution was decanted, and plates were washed 4 times with cold tap water for 30 minutes. After air-driying the plates overnight, the following day crystal violet was dissolved in 0.5ml of 10% acetic acid and absorbance of 200ul of sample was measured at 590nm using a TECAN Infinite M1000 Pro plate reader. For each treatment, the absorbance values for Aux-treated wells were normalized to the average absorbance of the untreated wells.

### Immunoblotting

Proteins were isolated from cells using a lysis buffer (50mM Tris-HCL, pH 8, 150mM NaCl, 0.1% Triton X-100, 0.5% sodium deoxycholate, 0.1% SDS, protease inhibitors (Millipore Sigma, P8340). 4-20% acrylamide gels (Bio-Rad, 4561096) were used to resolve the isolated proteins in running buffer (25mM Tris, 191mM Glycine, 0.1% SDS). PVDF membranes (Millipore Sigma, IPVH00010) were then used to transfer the proteins in transfer buffer (50mM Tris, 40mM Glycine, 0.005% SDS, 20% methanol). After blocking the membranes in blockinng buffer (50mM Tris, pH 7.5, 200mM NaCl, 0.1% Tween-20, 5% non-fat dry milk) for 1 hour at room temperature, diluted antibodies in TBSTM were added and incubated at 4°C overnight. To wash the membranes in the following day TBST (50mM Tris, pH 7.5, 200mM NaCl, 0.1% Tween-20) was used with 3 washes each for 10 minutes at room temperature. Next diluted secondary antibodies in TBST were added to the blots in room temperature for 1 hour. This was followed by three washes in TBST, each for 10 minutes at room temperature. Finally, the membranes were incubated with Immobilon Crescendo Western HRP substrate (Millipore Sigma, WBLUR0500) before imaging. Signal visualization was done using a Chemidoc Touch imaging system (Bio-Rad).

### Oligonucleotides and DNA fragments

Supplemental table 1.

### Antibodies

Supplemental table 2.

### Chemicals

Supplemental table 3.

---

**Supplemental Figure 1.**
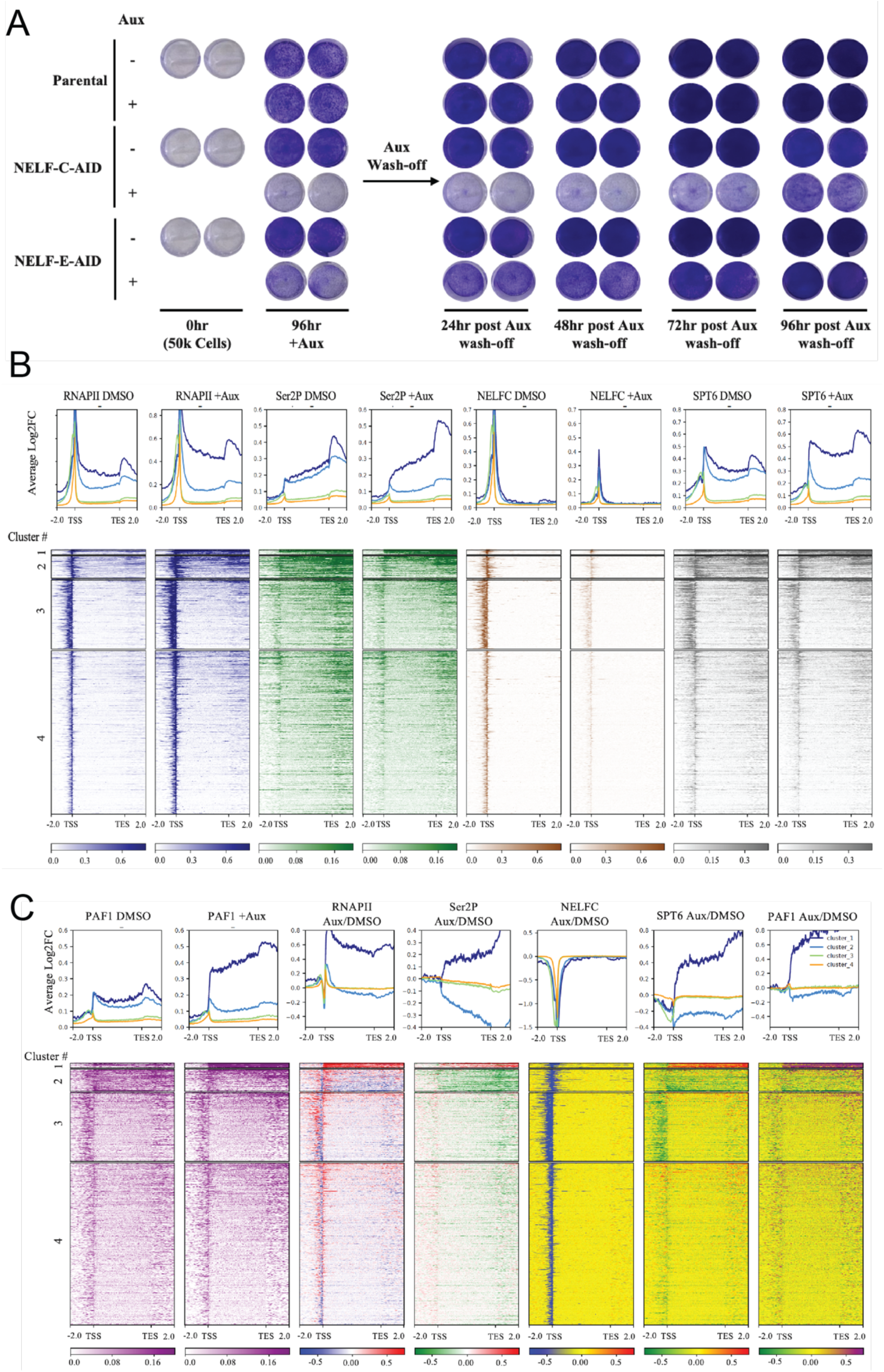
(A) Additional timepoints demonstrating dynamic reversal of the NELF depletion-induced growth defect observed in NELF-C-AID and NELF-E-AID but not parental DLD-1 cells, relevant to Figure 1A. Cells were seeded at 50k cells/well in 24-well plate at the beginning of the experiment (0hr), then treated with auxin (Aux, +) or DMSO control (-) for 96h prior to wash-off and culture in fresh media (24h, 48h, 72h and 96h timepoints shown), followed by fixation and crystal violet staining to visualize confluency. (B) Genome-wide metaplots (above) and heatmaps (below) of ChIP-seq signals (RNAPII, Ser2P, NELF-C, SPT6) in NELF-C-AID cells treated with either DMSO or auxin, relevant to Figure 1E. Genes have been divided into four clusters based on differential RNAPII occupancy. Cluster 1, n=111 genes. (C) Genome-wide metaplots (above) and heatmaps (below) of PAF1 ChIP-seq signal in NELF-C-AID cells treated with either DMSO or auxin (leftmost pair), relevant to Figure 1E, and of differential ChIP-seq signals (RNAPII, Ser2P, NELF-C, SPT6, PAF1) in auxin-treated relative to DMSO-treated NELF-C-AID cells, relevant to Figure 1E-F. Genes have been divided into four clusters as in (B).

**Supplemental Figure 2.**
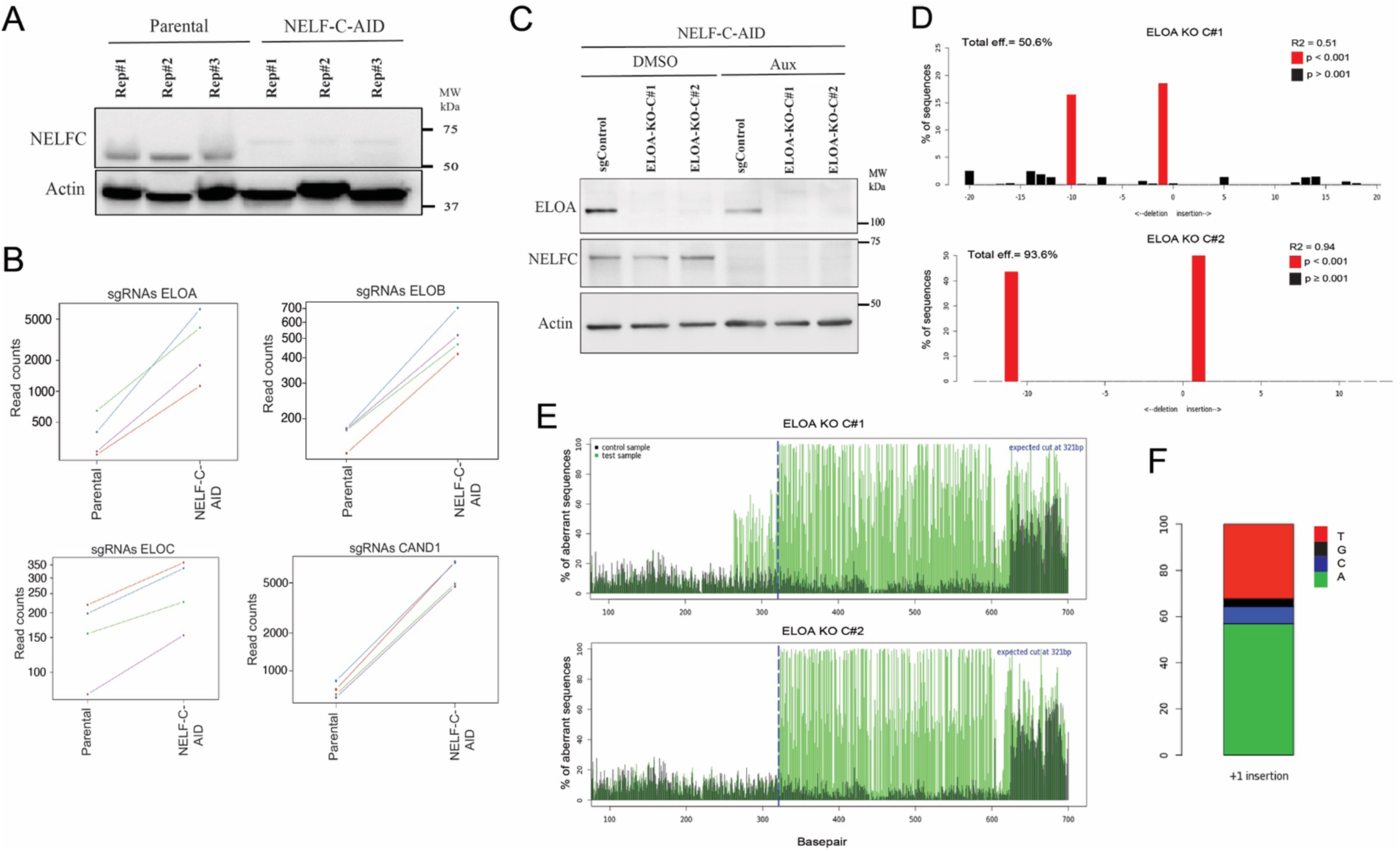
(A) Western blot for NELF-C, with blot for actin as a loading control, to confirm NELF-C depletion in auxin-treated NELF-C-AID but not parental DLD-1 cells. Three replicates are shown. (B) Read counts of individual sgRNA targeting the ELOA, ELOB, ELOC, and CAND1 genes in auxin-treated NELF-C-AID versus parental DLD-1 cells, illustrating relative enrichment of these sgRNA in the NELF-depleted condition. (C) Western blot showing clonal knockouts of ELOA in NELF-C-AID background. (D) sgRNA ELOA-KO confirmation as indel spectrum determined by TIDE, KO shown for two clones. (E) aberrant nucleotide signal of the KO sample (green) compared to that of the WT control (black). Blue dotted line indicates the expected cutting site. (F) The estimated composition of the inserted base for the +1-insertion modification in the ELOA-KO clone.

**Supplemental Figure 3.**
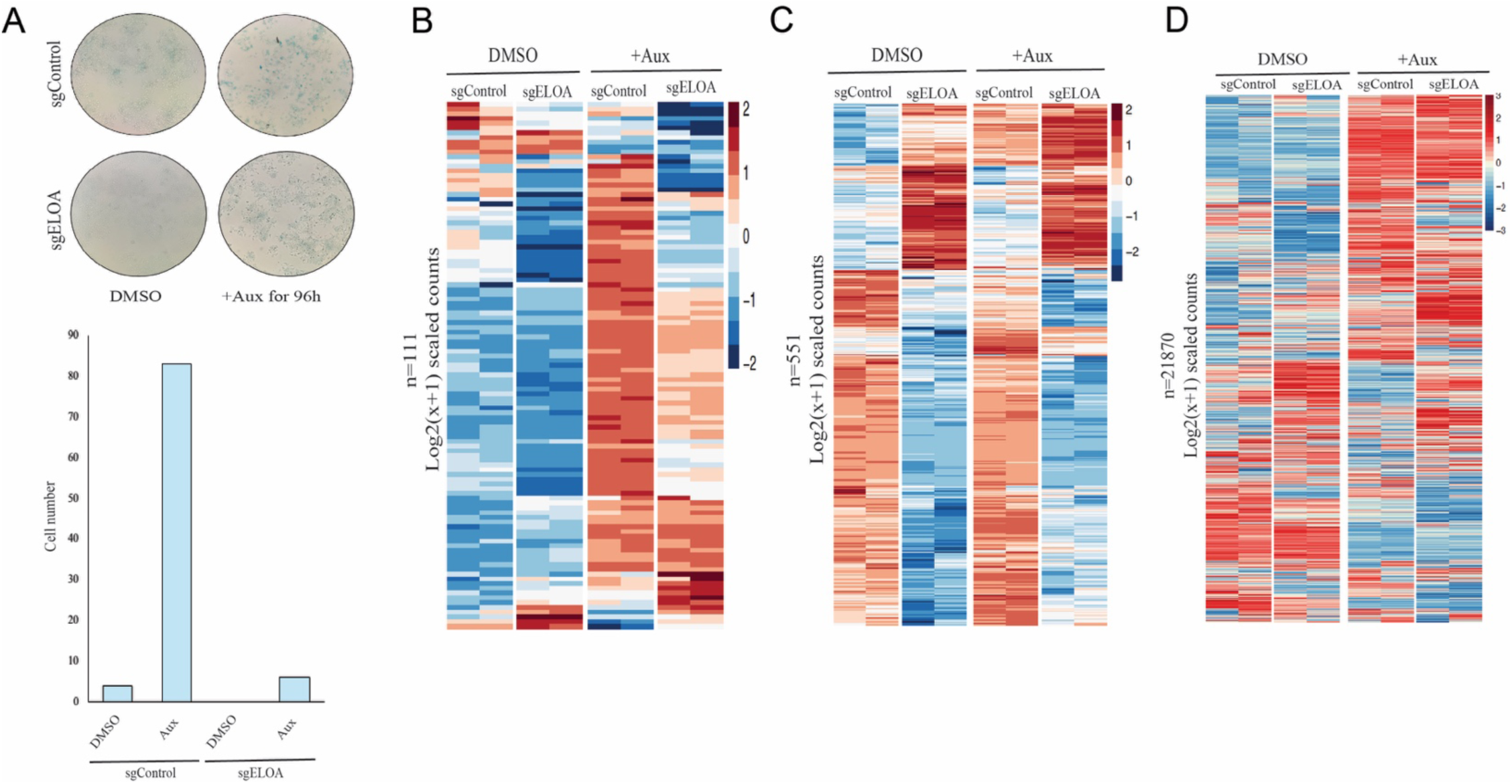
(A) β-galactosidase activity staining in NELF-C-AID cells when ELOA is knocked out. (B) Heatmap of gene expression levels (Log2-transformed, pseudocount-adjusted scaled RNA-seq read counts) in sgControl versus sgELOA NELF-C-AID cells treated with DMSO or auxin for 6h, for cluster 1 genes (the set of genes at which RNAPII is strongly released upon auxin treatment in NELF-C-AID cells, as shown in Fig. 1F, n=111 genes). (C) Heatmap of gene expression levels in sgControl versus sgELOA NELF-C-AID cells treated with DMSO or auxin for 6h, for the set of genes that are differentially expressed upon ELOA KO in untreated NELF-C-AID cells (n=551) (D) Heatmap of gene expression levels in sgControl versus sgELOA NELF-C-AID cells treated with DMSO or auxin for 6h, for all genes (n=21870).

**Supplemental Figure 4.**
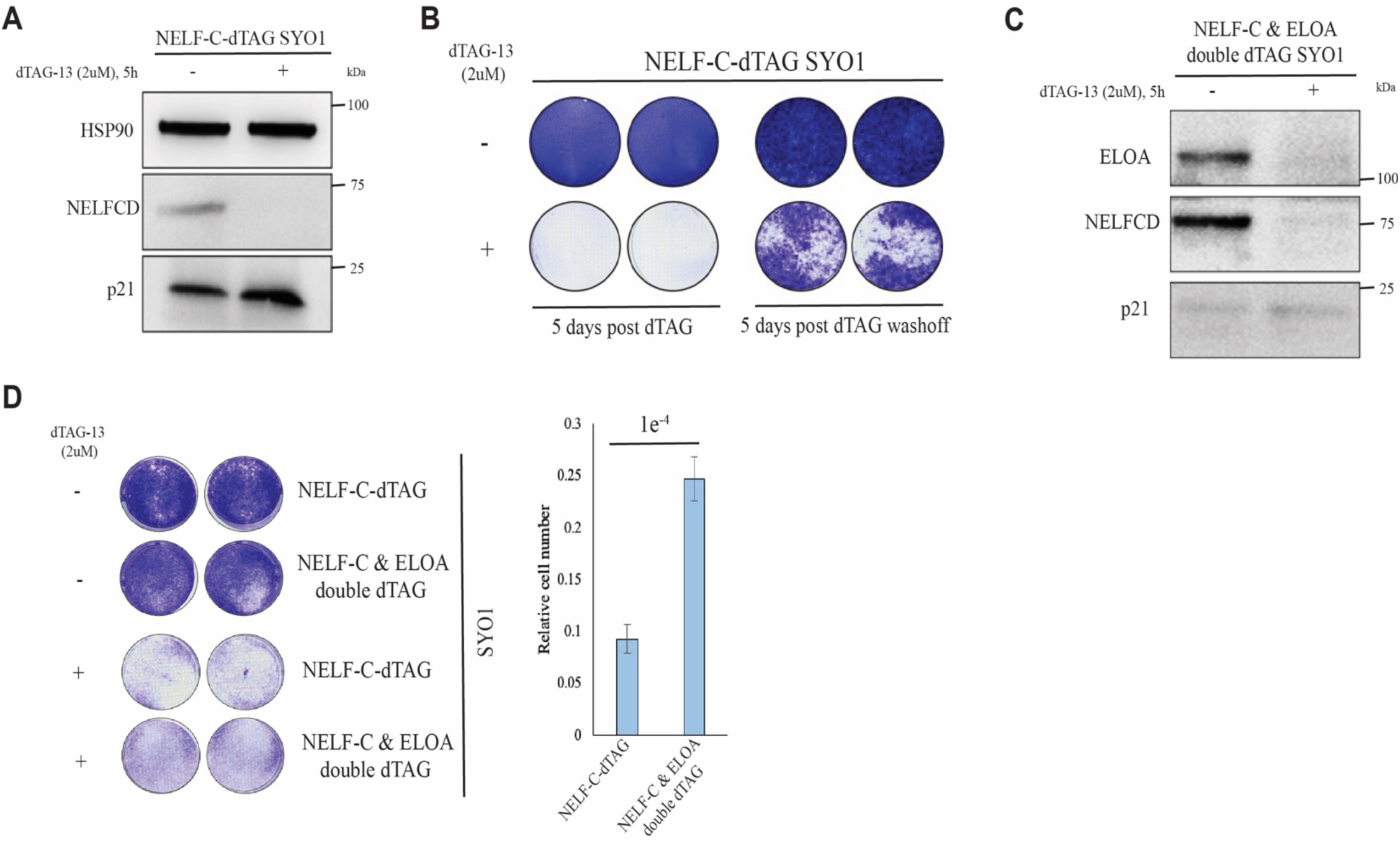
(A) Western blot analysis of NELFCD depletion in the NELF-C-dTAG SYO-1 cell line. Blots were probed for NELFCD and p21 with HSP90 as loading control. (B) Reversible growth defect in NELFCD-depleted cells. NELF-C-dTAG SYO-1 cells were treated with DMSO control (-) or dTAG-13 (2uM) for 96h prior to wash-off and culture in fresh media for 96h, followed by fixation and crystal violet staining to visualize confluency. (C) Western blot analysis of NELF-C and ELOA depletion upon dTAG-13 treatment (2uM, 5h) in the NELF-C and ELOA double dTAG cell line. Blots were probed for ELOA, NELFCD, and p21. (D) Rescue of the NELF depletion-induced growth defect observed in NELF-C-dTAG SYO-1 cells via additional depletion of ELOA in the NELF-C and ELOA double dTAG SYO-1 cell line. Cells were treated with DMSO or dTAG-13 (2uM) for 96h. The number of dTAG-treated relative to DMSO-treated cells is quantified at right.

**Supplemental Figure 5.**
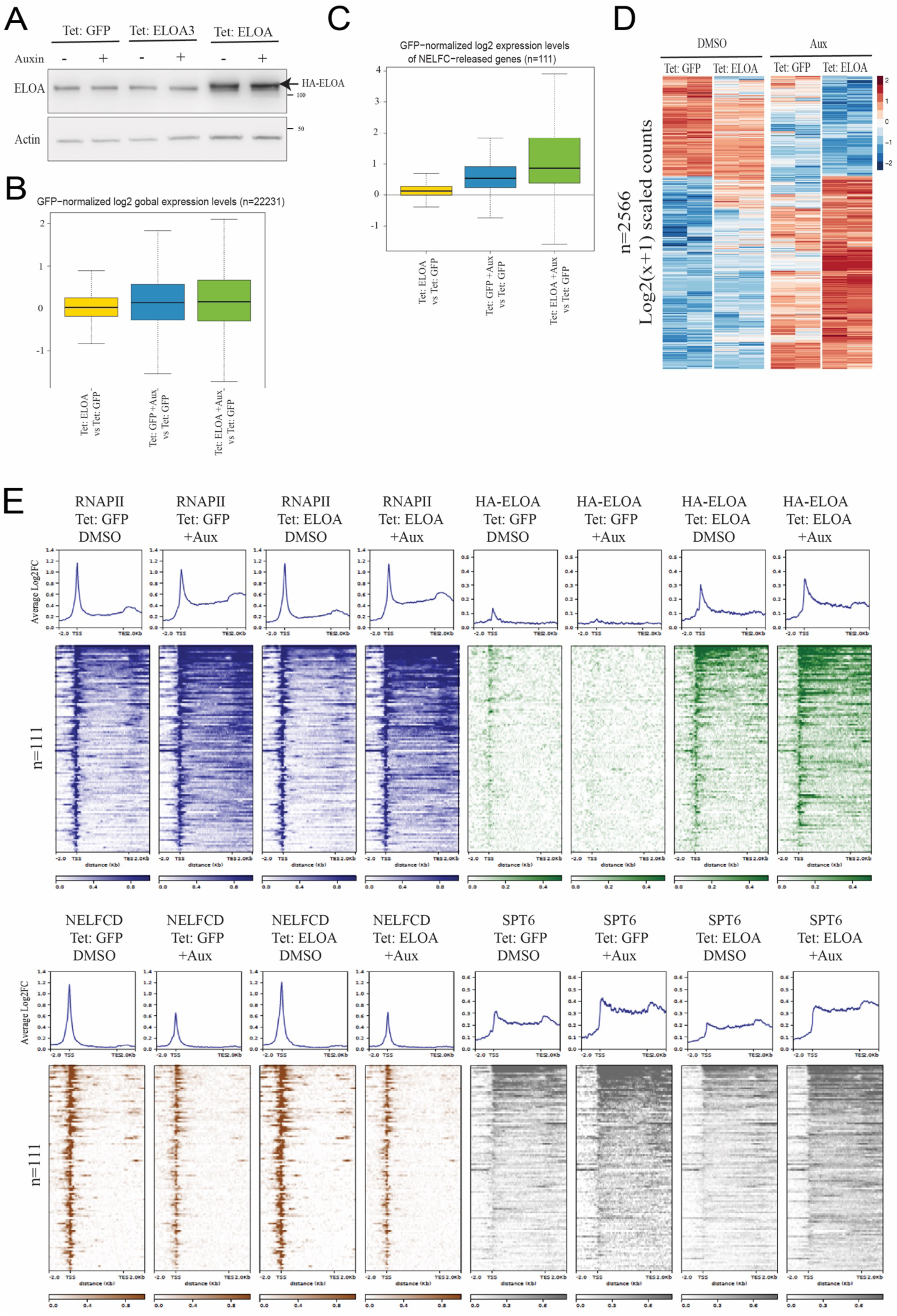
(A) Boxplots of global (n=22231) gene expression levels in untreated NELF-C-AID cells overexpressing ELOA (yellow) in comparison to auxin-treated NELF-C-AID cells overexpressing either GFP (blue) or ELOA (green). (B) Boxplots of gene expression levels as in (A), for cluster 1 genes (n=111), relevant to Fig. 3E. (C) Heatmap of gene expression levels (Log2-transformed, pseudocount-adjusted scaled RNA-seq read counts) in NELF-C-AID cells overexpressing GFP or ELOA and treated with DMSO or auxin for 6h, for the set of genes that are differentially expressed in the auxin-treated cells overexpressing ELOA relative to DMSO-treated cells overexpressing GFP (n=2566). (D) Genome-wide metaplots (above) and heatmaps (below) of ChIP-seq signals for RNAPII (blue), ELOA (green), NELF-C (red), and PAF1 (purple) in NELF-C-AID cells overexpressing GFP or ELOA and treated with either DMSO or auxin, relevant to Figure 3H.

**Supplemental Figure 6.**
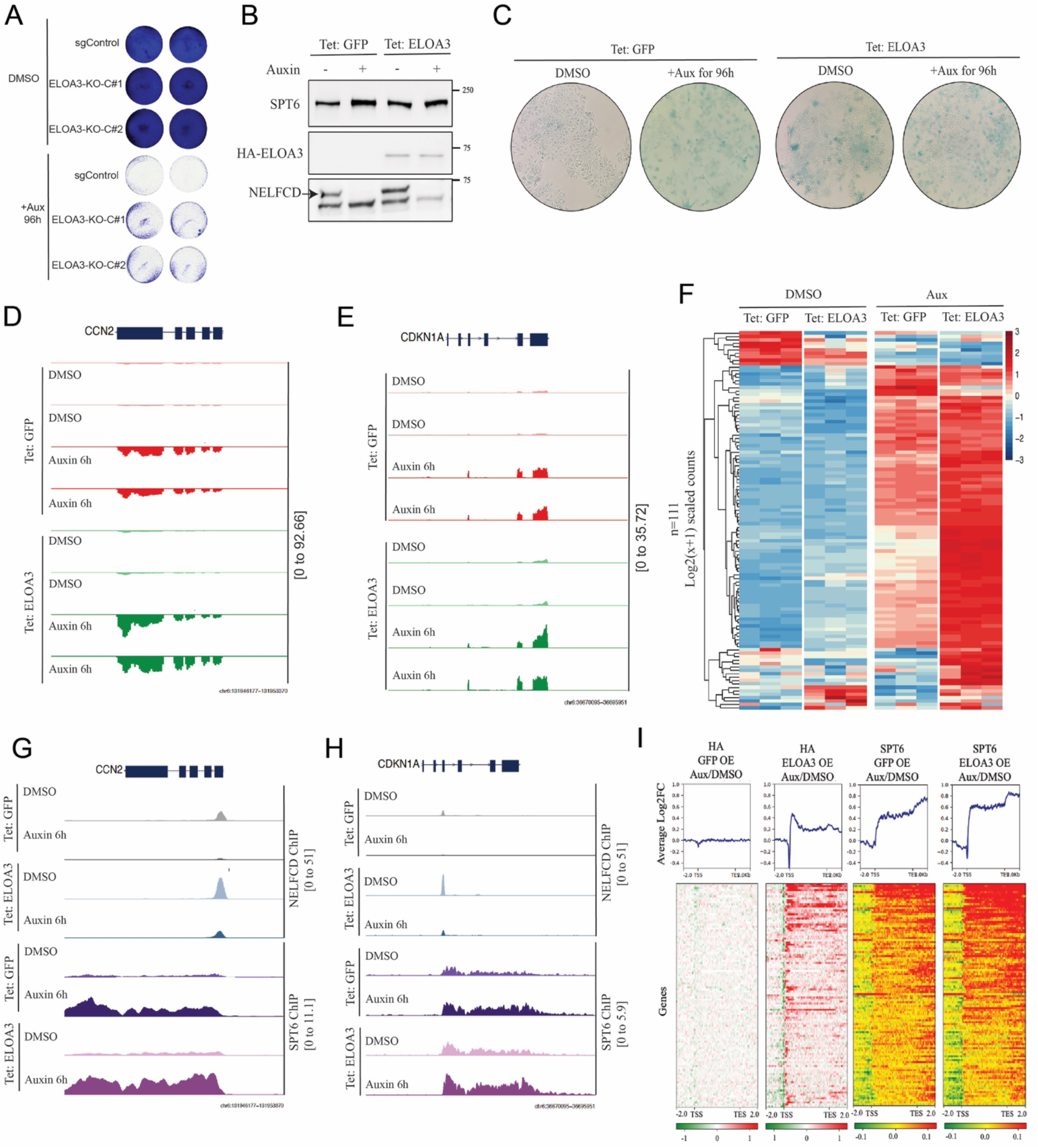
(A) NELF depletion-induced growth defect in NELF-C-AID cells can be rescued with ELOA3 clonal knockouts using two distinct ELOA3-targeting sgRNA (ELOA3-KO-C#1 and #2), with a nontargeting sgRNA as control (sgControl). Auxin (+Aux) or DMSO control (-Aux) treatment was done for 8 days with fresh auxin supplemented daily. The number of auxin-treated relative to DMSO-treated cells is quantified at right. (B) Western blots confirmation of generation of cell lines stably overexpressing N-terminal FLAG and HA-tagged ELOA3 on top of NELF-C-AID. Cells were treated with DMSO (-) or auxin (+) for 6h and Dox for 24h. (C) Staining for senescence-associated β-galactosidase activity in NELF-C-AID cells stably expressing GFP or ELOA3 and treated with DMSO or auxin for 96h. (D-E) Genome browser track visualizations of RNA-seq signal (two replicates shown) at the senescence-associated genes CCN2 (D) and CDKN1A (E) in NELF-C-AID cells stably expressing GFP or ELOA3 and treated with DMSO or auxin for 6h. (F) Hierarchical clustering heatmap of Log2-transformed, pseudocount-adjusted RNA-seq reads in NELF-C-AID cells stably expressing GFP or ELOA3 and treated with DMSO or auxin for 6h, for cluster 1 genes (the set of genes at which RNAPII is strongly released upon auxin treatment in NELF-C-AID cells as shown in Fig. 1F, n=111 genes). (G-H) Genome browser track visualizations of NELFC and SPT6 ChIP-seq signals at CCN2 (G) and CDKN1A (H) in NELF-C-AID cells stably expressing GFP or ELOA3 and treated with DMSO or auxin for 6h. (I) Metaplots of Log2-transformed fold change (Log2FC, above) and heatmaps (below) of differential ELOA3 (HA tag) and SPT6 ChIP-seq signal at the same set of genes as in (F), in auxin-treated relative to DMSO-treated NELF-C-AID cells stably expressing GFP or ELOA3 (FLAG and HA-tagged). TSS= transcription start site; TES=transcription end site. n=111 genes.

**Supplemental Figure 7.**
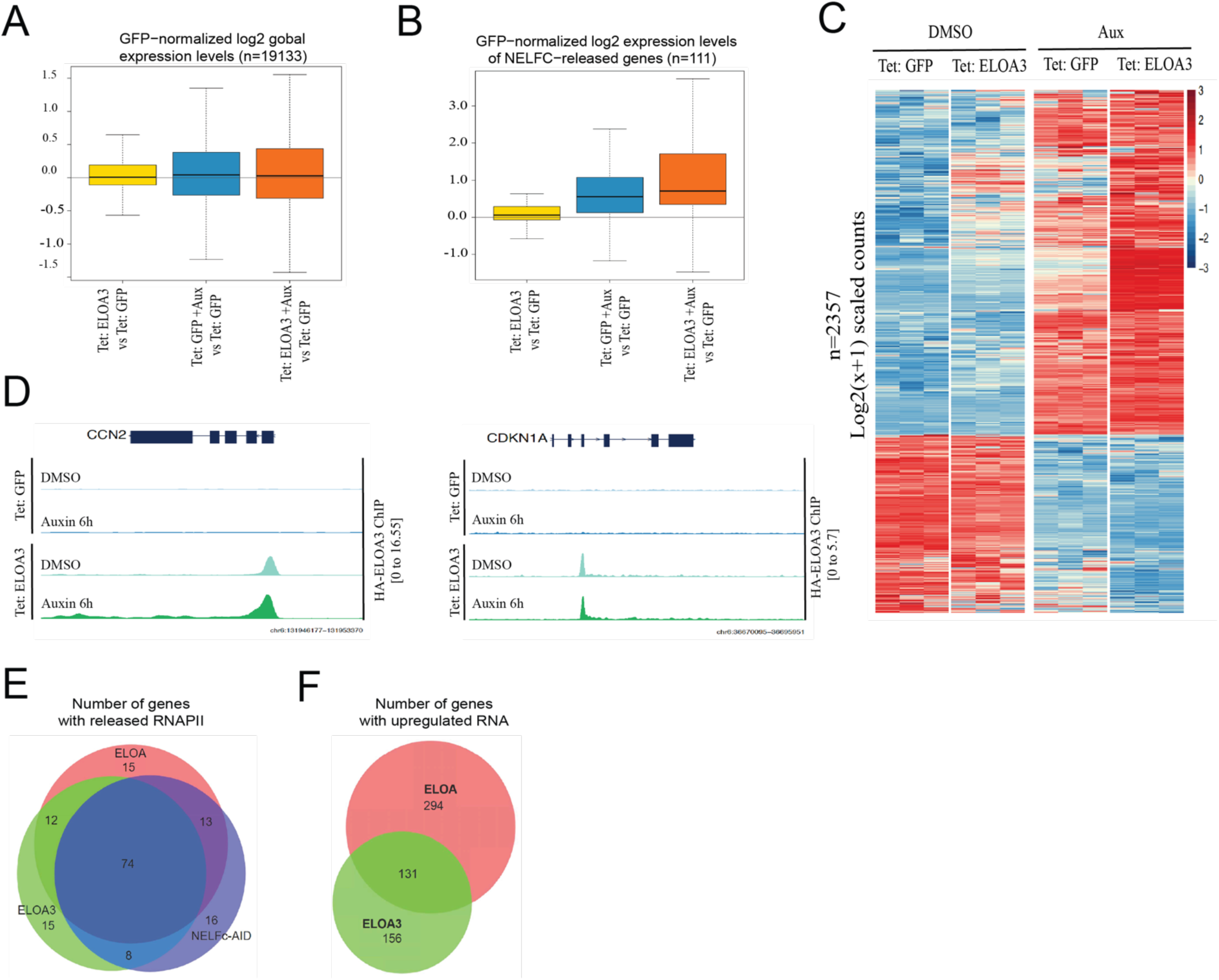
(A) Boxplots of global (n=19133) gene expression levels in untreated NELF-C-AID cells overexpressing ELOA3 (yellow) in comparison to auxin-treated NELF-C-AID cells overexpressing either GFP (blue) or ELOA (orange). (B) Boxplots of gene expression levels for cluster 1 genes (n=111). (C) Heatmap of gene expression levels (Log2-transformed, pseudocount-adjusted scaled RNA-seq read counts) in NELF-C-AID cells stably expressing GFP or ELOA3 and treated with DMSO or auxin for 6h, for the set of all genes at which NELF-C depletion-induced expression changes were amplified by ELOA KO (n=2357). (D) Genome browser track visualizations of RNAPII and ELOA3 ChIP-seq signals at the senescence-associated genes CCN2 and CDKN1A in NELF-C-AID cells stably expressing GFP or ELOA3 and treated with DMSO or auxin for 6h. (E) Venn diagram showing the number of genes with released RNAPII when comparing Auxin to DMSO treatments (for NELFC depletion) and simultaneously overexpressing Tet: GFP, Tet: ELOA or Tet:ELOA3. (F) Venn diagram in which number of genes upregulated at the mRNA are shown for conditions with NELFC depletion and overexpression of ELOA or ELOA3. These gene numbers are normalized to a condition for which NELFC is depleted and GFP is overexpressed.

**Supplemental Table S1.**
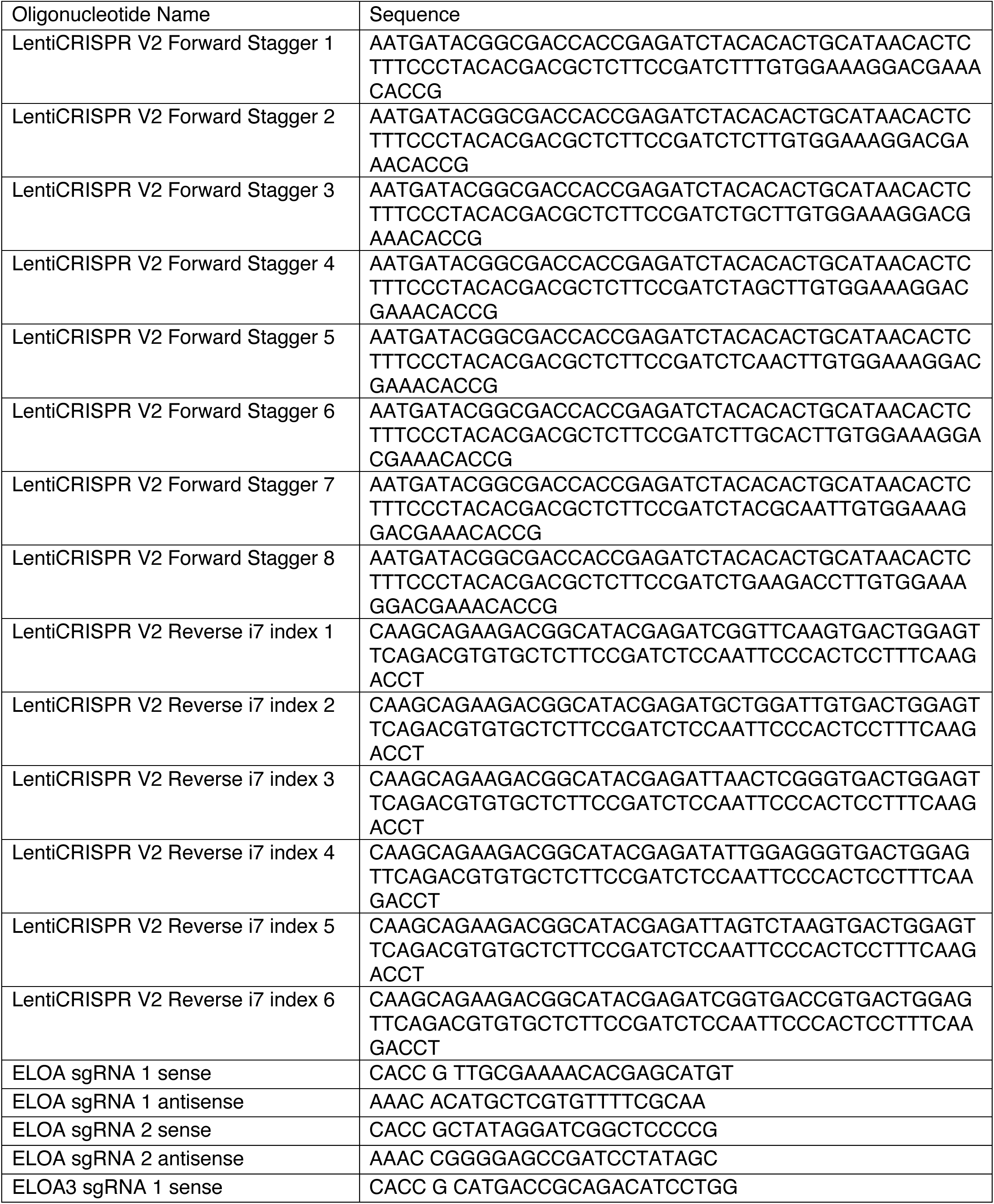
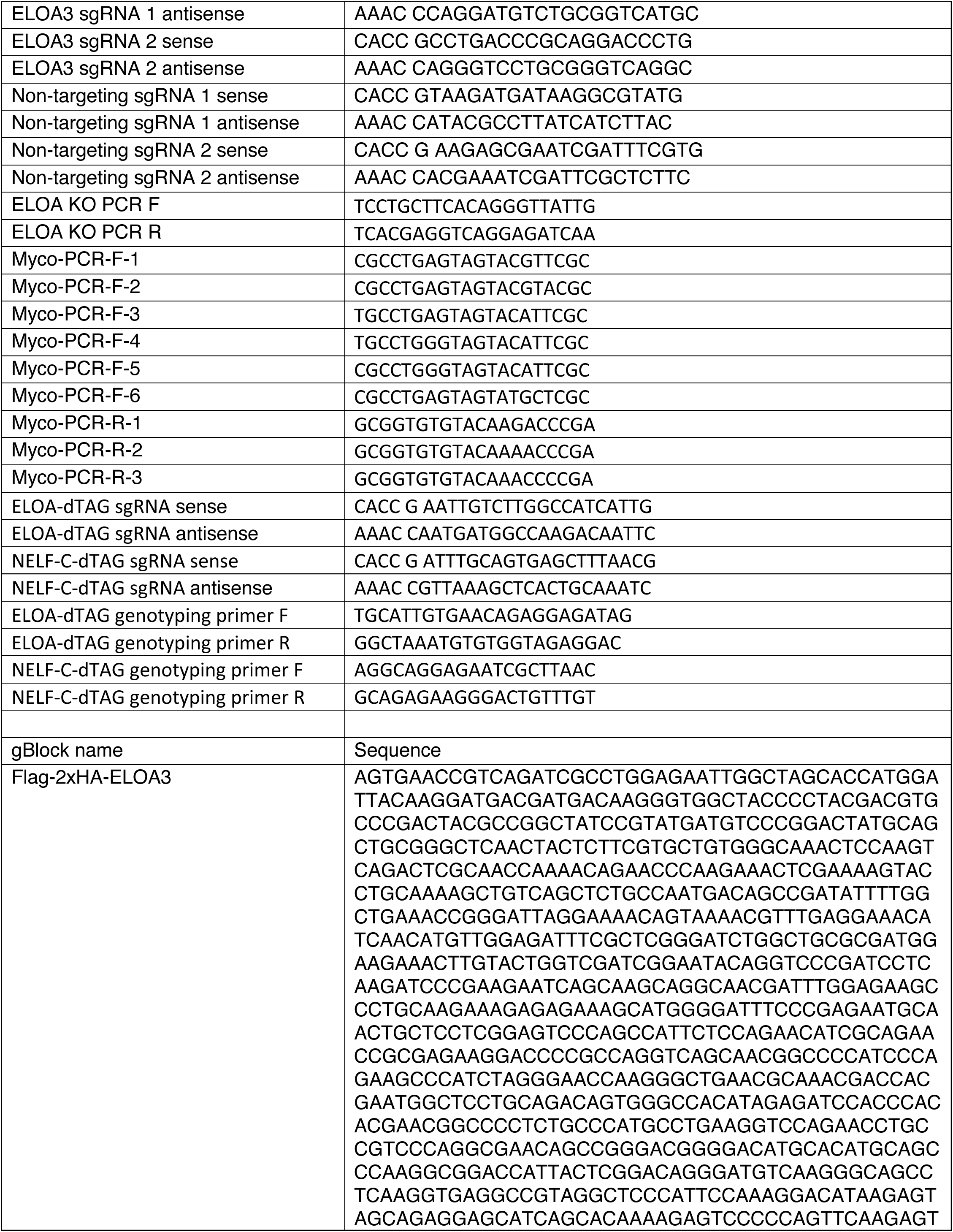
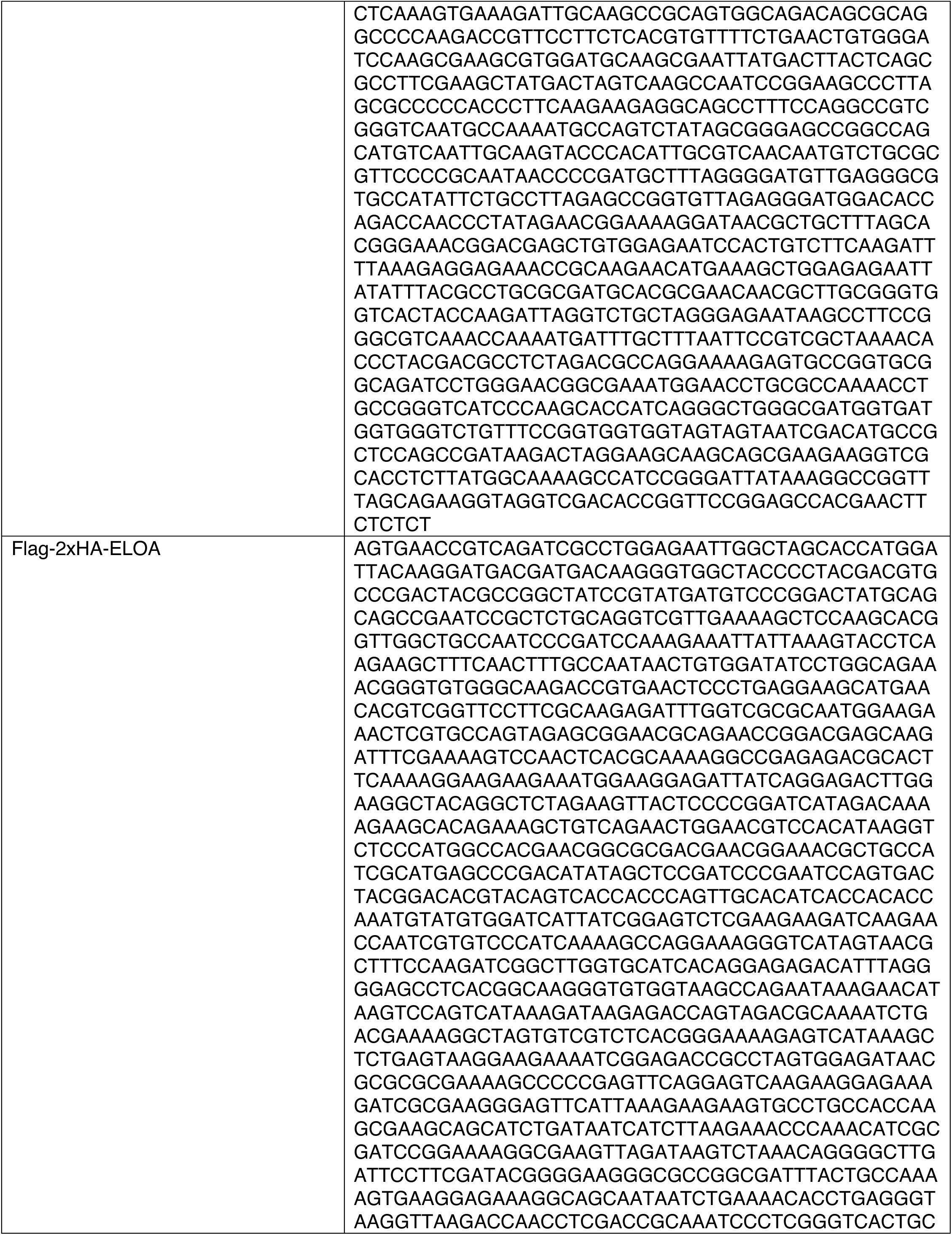
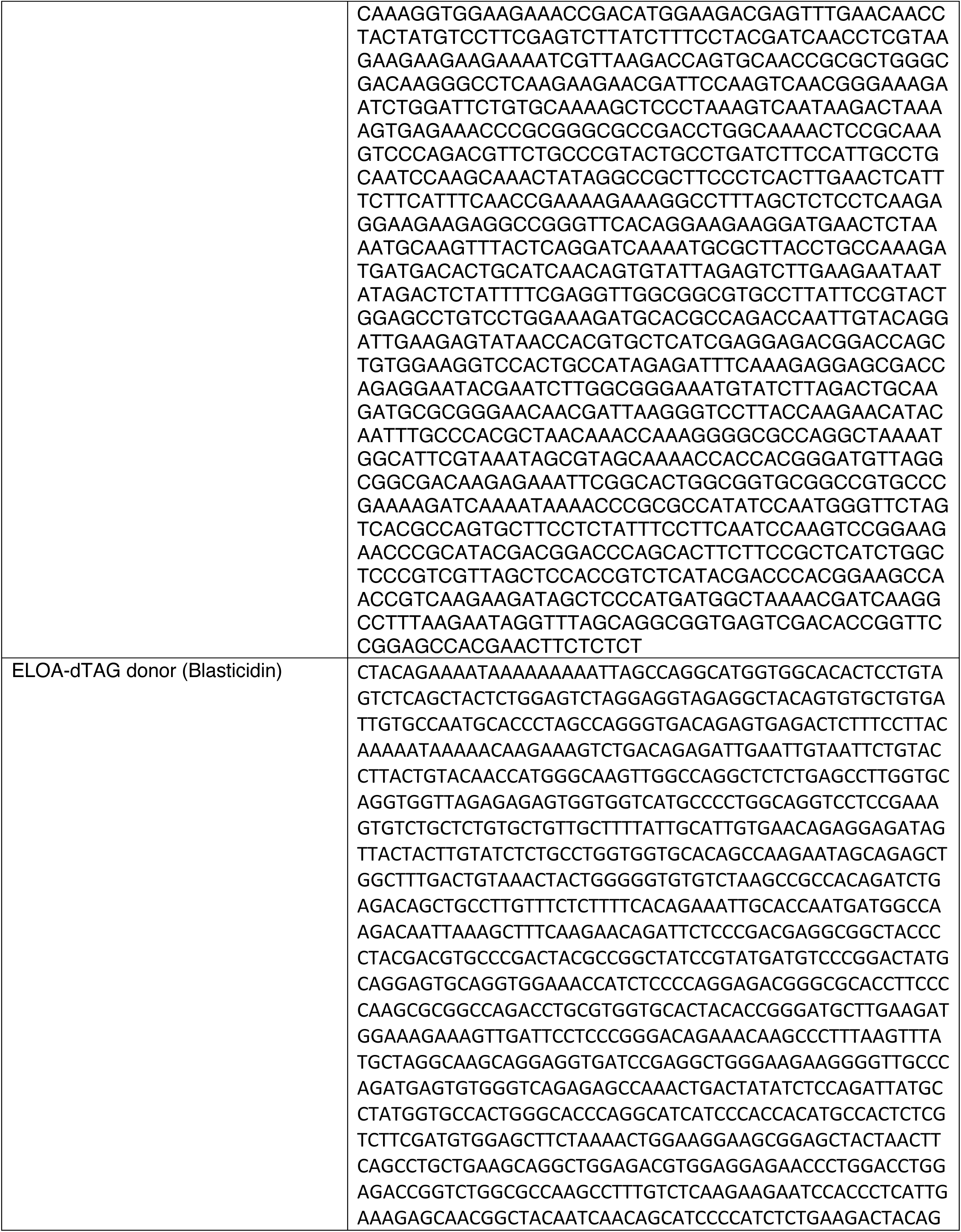
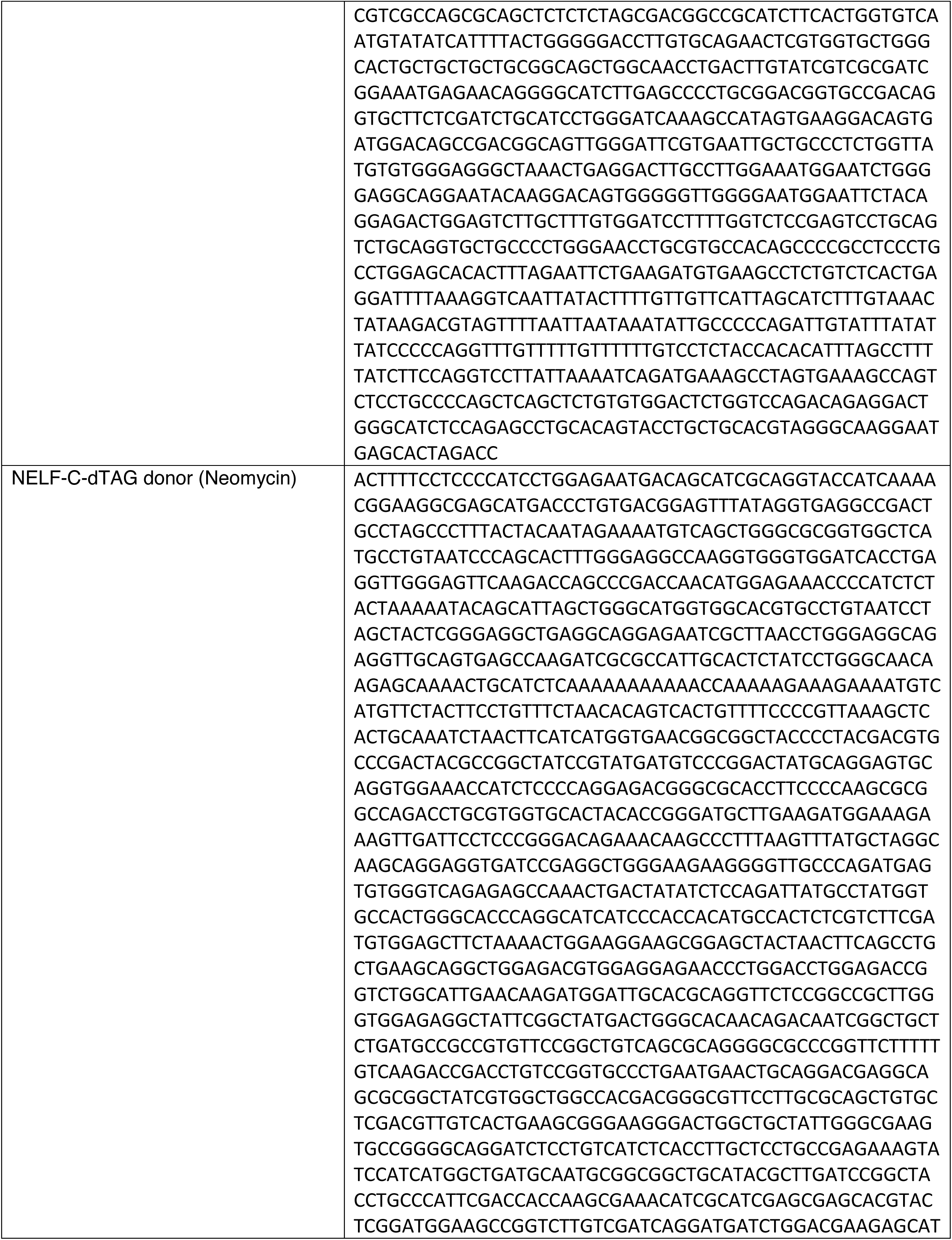
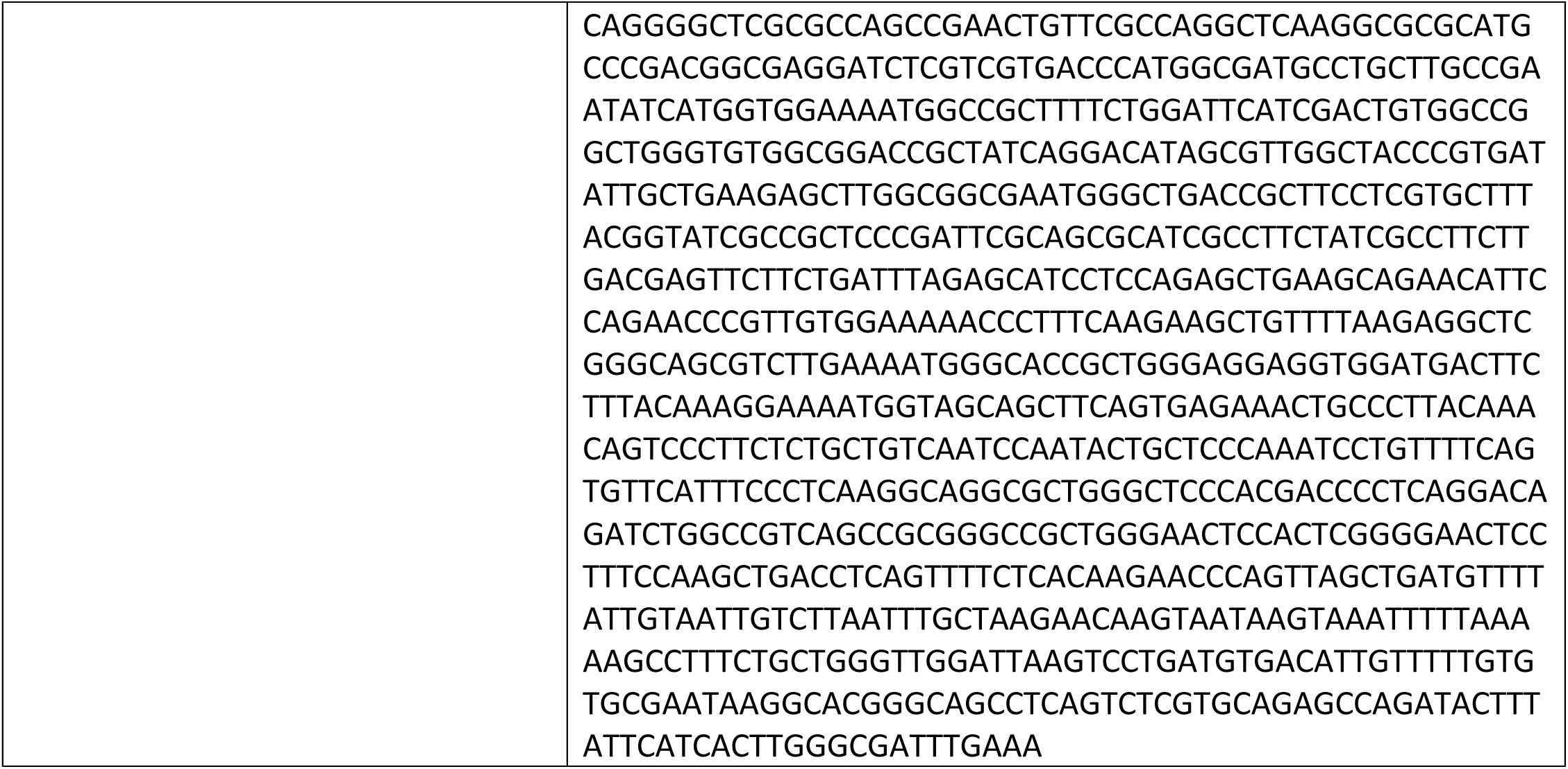
Oligonucleotides and synthetic DNA.

**Supplemental Table S2.** Antibodies.

| **<u>Name</u>** | **<u>Supplier</u>** | **<u>Catalog #</u>** | **<u>Immunoblot</u>**<br>**<u>ting</u>** | **<u>ChIP</u>** | **<u>CUT&RUN</u>** |
| --- | --- | --- | --- | --- | --- |
| HA-tag | Cell Signaling Technology | 3724S | 1:2,000 | 10ul / 1mg chromatin | N/A |
| GFP | Santa Cruz | sc-9996 | 1:1,000 | N/A | N/A |
| Actin | Santa Cruz | sc-13119 | 1:10,000 | N/A | N/A |
| p21/CDKN1A | Cell Signaling Technology | 2947S | 1:1,000 | N/A | N/A |
| POLR2A (Rpb1) NTD | Cell Signaling Technology |  | 1:1,000 | 10ul / 1mg chromatin | N/A |
| POLR2A (Rpb1) CTD | Cell Signaling Technology | 14958S | 1:1,000 | N/A |  |
| SPT6 | Cell Signaling Technology | 15616S | 1:1,000 | 10ul / 1mg chromatin | N/A |
| PAF1 | Cell Signaling Technology | 12883S | N/A | 10ul / 1mg chromatin | N/A |
| POLR2A (Rpb1) pSer2 | Cell Signaling Technology | 13499S | N/A | 10ul / 1mg chromatin | 1:50 |
| CCN2 | Cell Signaling Technology | 86641S | 1:1,000 | N/A | N/A |
| NELFCD | Cell Signaling Technology | 12265S | 1:1,000 | 10ul / 1mg chromatin | N/A |
| ELOA | Bethyl Laboratories | A300-942A | 1:1,000 | 10ul / 1mg chromatin | N/A |
| PARP | Cell Signaling Technology | 5625S | 1:1,000 | N/A | N/A |
| Anti-Rabbit HRP | Millipore Sigma | NA934V | 1:2,000 | N/A | N/A |
| Anti-Mouse HRP | Millipore Sigma | NA931V | 1:2,000 | N/A | N/A |

N/A: Not Applicable to this study

**Supplemental Table S3.** Chemicals.

| ***<u>Chemical name</u>*** | ***<u>Supplier</u>*** | ***<u>Catalog #</u>*** |
| --- | --- | --- |
| X-Gal | Thermo Fisher Scientific | R0404 |
| Indole-3-acetic acid sodium salt | Sigma-Aldrich | I5148-2G |
| Doxycycline hyclate | Millipore Sigma | D9891-1G |
| Puromycin | Millipore Sigma | P8833-25MG |
| Hygromycin B | Thermo Fisher Scientific | 10687010 |
| dTAG-13 | Tocris | 6605 |
| Geneticin Selective Antibiotic (G418 Sulfate) | Thermo Fisher Scientific | 10131035 |

## References

1. Y. Aoi, A. Shilatifard, Transcriptional elongation control in developmental gene expression, aging, and disease. Mol Cell 83, 3972–3999 (2023).

2. F. Chen, X. Gao, A. Shilatifard, Stably paused genes revealed through inhibition of transcription initiation by the TFIIH inhibitor triptolide. Genes Dev 29, 39–47 (2015).

3. L. Core, K. Adelman, Promoter-proximal pausing of RNA polymerase II: a nexus of gene regulation. Genes Dev 33, 960–982 (2019).

4. G. W. Muse et al., RNA polymerase is poised for activation across the genome. Nat Genet 39, 1507–1511 (2007).

5. B. Stadelmayer et al., Integrator complex regulates NELF-mediated RNA polymerase II pause/release and processivity at coding genes. Nat Commun 5, 5531 (2014).

6. Y. Aoi et al., SPT5 regulates RNA polymerase II stability via Cullin 3-ARMC5 recognition. Sci Adv 11, eadt5885 (2025).

7. Y. Aoi et al., SPT5 stabilization of promoter-proximal RNA polymerase II. Mol Cell 81, 4413–4424 e4415 (2021).

8. Y. Yamaguchi et al., NELF, a multisubunit complex containing RD, cooperates with DSIF to repress RNA polymerase II elongation. Cell 97, 41–51 (1999).

9. C. Evrin et al., Spt5 histone binding activity preserves chromatin during transcription by RNA polymerase II. Embo j 41, e109783 (2022).

10. F. X. Chen, E. R. Smith, A. Shilatifard, Born to run: control of transcription elongation by RNA polymerase II. Nat Rev Mol Cell Biol 19, 464–478 (2018).

11. Z. Luo, C. Lin, A. Shilatifard, The super elongation complex (SEC) family in transcriptional control. Nat Rev Mol Cell Biol 13, 543–547 (2012).

12. S. M. Vos et al., Structure of activated transcription complex Pol II-DSIF-PAF-SPT6. Nature 560, 607–612 (2018).

13. S. M. Vos, L. Farnung, H. Urlaub, P. Cramer, Structure of paused transcription complex Pol II-DSIF-NELF. Nature 560, 601–606 (2018).

14. T. Narita et al., Human transcription elongation factor NELF: identification of novel subunits and reconstitution of the functionally active complex. Mol Cell Biol 23, 1863–1873 (2003).

15. Y. Aoi et al., NELF Regulates a Promoter-Proximal Step Distinct from RNA Pol II Pause-Release. Mol Cell 78, 261–274 e265 (2020).

16. Y. Aoi et al., SPT6 functions in transcriptional pause/release via PAF1C recruitment. Mol Cell 82, 3412–3423 e3415 (2022).

17. S. M. Yoh, H. Cho, L. Pickle, R. M. Evans, K. A. Jones, The Spt6 SH2 domain binds Ser2-P RNAPII to direct Iws1-dependent mRNA splicing and export. Genes Dev 21, 160–174 (2007).

18. M. B. Ardehali et al., Spt6 enhances the elongation rate of RNA polymerase II in vivo. EMBO J 28, 1067–1077 (2009).

19. T. Aso, W. S. Lane, J. W. Conaway, R. C. Conaway, Elongin (SIII): a multisubunit regulator of elongation by RNA polymerase II. Science 269, 1439–1443 (1995).

20. J. N. Bradsher, K. W. Jackson, R. C. Conaway, J. W. Conaway, RNA polymerase II transcription factor SIII. I. Identification, purification, and properties. J Biol Chem 268, 25587–25593 (1993).

21. J. N. Bradsher, S. Tan, H. J. McLaury, J. W. Conaway, R. C. Conaway, RNA polymerase II transcription factor SIII. II. Functional properties and role in RNA chain elongation. J Biol Chem 268, 25594–25603 (1993).

22. M. B. Ardehali, M. Damle, C. Perea-Resa, M. D. Blower, R. E. Kingston, Elongin A associates with actively transcribed genes and modulates enhancer RNA levels with limited impact on transcription elongation rate in vivo. J Biol Chem 296, 100202 (2021).

23. Y. Wang, L. Hou, M. B. Ardehali, R. E. Kingston, B. D. Dynlacht, Elongin A regulates transcription in vivo through enhanced RNA polymerase processivity. J Biol Chem 296, 100170 (2021).

24. M. Gerber et al., Regulation of heat shock gene expression by RNA polymerase II elongation factor, Elongin A. J Biol Chem 280, 4017–4020 (2005).

25. J. Kawauchi et al., Transcriptional properties of mammalian elongin A and its role in stress response. J Biol Chem 288, 24302–24315 (2013).

26. Y. Chen et al., Structure of the transcribing RNA polymerase II-Elongin complex. Nat Struct Mol Biol 30, 1925–1935 (2023).

27. J. C. Weems et al., Assembly of the Elongin A Ubiquitin Ligase Is Regulated by Genotoxic and Other Stresses. J Biol Chem 290, 15030–15041 (2015).

28. T. Yasukawa et al., Mammalian Elongin A complex mediates DNA-damage-induced ubiquitylation and degradation of Rpb1. EMBO J 27, 3256–3266 (2008).

29. I. Olan, M. Narita, Senescence: An Identity Crisis Originating from Deep Within the Nucleus. Annu Rev Cell Dev Biol 38, 219–239 (2022).

30. D. Saul et al., A new gene set identifies senescent cells and predicts senescence-associated pathways across tissues. Nat Commun 13, 4827 (2022).

31. H. P. Gala et al., A transcriptionally repressed quiescence program is associated with paused RNA polymerase II and is poised for cell cycle re-entry. J Cell Sci 135, (2022).

32. A. Gyenis et al., Genome-wide RNA polymerase stalling shapes the transcriptome during aging. Nat Genet 55, 268–279 (2023).

33. K. E. C. Blokland, S. D. Pouwels, M. Schuliga, D. A. Knight, J. K. Burgess, Regulation of cellular senescence by extracellular matrix during chronic fibrotic diseases. Clin Sci (Lond) 134, 2681–2706 (2020).

34. C. Capparelli et al., CTGF drives autophagy, glycolysis and senescence in cancer-associated fibroblasts via HIF1 activation, metabolically promoting tumor growth. Cell Cycle 11, 2272–2284 (2012).

35. J. I. Jun, L. F. Lau, CCN2 induces cellular senescence in fibroblasts. J Cell Commun Signal 11, 15–23 (2017).

36. F. X. Chen et al., PAF1 regulation of promoter-proximal pause release via enhancer activation. Science 357, 1294–1298 (2017).

37. N. L. Bray, H. Pimentel, P. Melsted, L. Pachter, Near-optimal probabilistic RNA-seq quantification. Nat Biotechnol 34, 525–527 (2016).

38. F. J. Pardo-Palacios et al., SQANTI3: curation of long-read transcriptomes for accurate identification of known and novel isoforms. Nat Methods 21, 793–797 (2024).

39. M. Apostolides et al., Accurate isoform quantification by joint short- and long-read RNA-sequencing. bioRxiv, (2024).

40. A. Bakulin, N. B. Teyssier, M. Kampmann, M. Khoroshkin, H. Goodarzi, pyPAGE: A framework for Addressing biases in gene-set enrichment analysis-A case study on Alzheimer’s disease. PLoS Comput Biol 20, e1012346 (2024).

41. M. J. McBride et al., The SS18-SSX Fusion Oncoprotein Hijacks BAF Complex Targeting and Function to Drive Synovial Sarcoma. Cancer Cell 33, 1128–1141 e1127 (2018).

42. M. A. J. Morgan et al., ELOA3: A primate-specific RNA polymerase II elongation factor encoded by a tandem repeat gene cluster. Sci Adv 9, eadj1261 (2023).

43. L. Davidson, L. Muniz, S. West, 3’ end formation of pre-mRNA and phosphorylation of Ser2 on the RNA polymerase II CTD are reciprocally coupled in human cells. Genes Dev 28, 342–356 (2014).

44. T. Tufan et al., Rapid unleashing of macrophage efferocytic capacity via transcriptional pause release. Nature 628, 408–415 (2024).

45. L. Yu et al., Negative elongation factor complex enables macrophage inflammatory responses by controlling anti-inflammatory gene expression. Nat Commun 11, 2286 (2020).

46. K. Fujimaki et al., Graded regulation of cellular quiescence depth between proliferation and senescence by a lysosomal dimmer switch. Proc Natl Acad Sci U S A 116, 22624–22634 (2019).

47. H. M. Ashraf, B. Fernandez, S. L. Spencer, The intensities of canonical senescence biomarkers integrate the duration of cell-cycle withdrawal. Nat Commun 14, 4527 (2023).

48. Z. K. Ngian, W. Q. Lin, C. T. Ong, NELF-A controls Drosophila healthspan by regulating heat-shock protein-mediated cellular protection and heterochromatin maintenance. Aging Cell 20, e13348 (2021).

49. E. Yang et al., Decay rates of human mRNAs: correlation with functional characteristics and sequence attributes. Genome Res 13, 1863–1872 (2003).

50. L. W. Harries et al., Human aging is characterized by focused changes in gene expression and deregulation of alternative splicing. Aging Cell 10, 868–878 (2011).

51. A. C. Holly et al., Changes in splicing factor expression are associated with advancing age in man. Mech Ageing Dev 134, 356–366 (2013).

52. B. P. Lee et al., Changes in the expression of splicing factor transcripts and variations in alternative splicing are associated with lifespan in mice and humans. Aging Cell 15, 903–913 (2016).

53. S. Adusumalli, Z. K. Ngian, W. Q. Lin, T. Benoukraf, C. T. Ong, Increased intron retention is a post-transcriptional signature associated with progressive aging and Alzheimer’s disease. Aging Cell 18, e12928 (2019).

54. C. Heintz et al., Splicing factor 1 modulates dietary restriction and TORC1 pathway longevity in C. elegans. Nature 541, 102–106 (2017).

55. S. S. Tabrez, R. D. Sharma, V. Jain, A. A. Siddiqui, A. Mukhopadhyay, Differential alternative splicing coupled to nonsense-mediated decay of mRNA ensures dietary restriction-induced longevity. Nat Commun 8, 306 (2017).

56. H. Shenasa, D. L. Bentley, Pre-mRNA splicing and its cotranscriptional connections. Trends Genet 39, 672–685 (2023).

57. K. Miyata et al., Induction of apoptosis and cellular senescence in mice lacking transcription elongation factor, Elongin A. Cell Death Differ 14, 716–726 (2007).

58. Y. Katz et al., Quantitative visualization of alternative exon expression from RNA-seq data. Bioinformatics 31, 2400–2402 (2015).

59. J. G. Doench et al., Optimized sgRNA design to maximize activity and minimize off-target effects of CRISPR-Cas9. Nat Biotechnol 34, 184–191 (2016).

60. W. Li et al., MAGeCK enables robust identification of essential genes from genome-scale CRISPR/Cas9 knockout screens. Genome Biol 15, 554 (2014).

61. T. I. Lee, S. E. Johnstone, R. A. Young, Chromatin immunoprecipitation and microarray-based analysis of protein location. Nat Protoc 1, 729–748 (2006).

62. B. Langmead, C. Trapnell, M. Pop, S. L. Salzberg, Ultrafast and memory-efficient alignment of short DNA sequences to the human genome. Genome Biol 10, R25 (2009).

63. Y. Zhang et al., Model-based analysis of ChIP-Seq (MACS). Genome Biol 9, R137 (2008).

64. F. Ramirez, F. Dundar, S. Diehl, B. A. Gruning, T. Manke, deepTools: a flexible platform for exploring deep-sequencing data. Nucleic Acids Res 42, W187–191 (2014).

65. U. Raudvere et al., g:Profiler: a web server for functional enrichment analysis and conversions of gene lists (2019 update). Nucleic Acids Res 47, W191–W198 (2019).

66. A. M. Bolger, M. Lohse, B. Usadel, Trimmomatic: a flexible trimmer for Illumina sequence data. Bioinformatics 30, 2114–2120 (2014).

67. A. Dobin et al., STAR: ultrafast universal RNA-seq aligner. Bioinformatics 29, 15–21 (2013).

68. S. Anders, P. T. Pyl, W. Huber, HTSeq--a Python framework to work with high-throughput sequencing data. Bioinformatics 31, 166–169 (2015).

69. D. Risso, J. Ngai, T. P. Speed, S. Dudoit, Normalization of RNA-seq data using factor analysis of control genes or samples. Nat Biotechnol 32, 896–902 (2014).

70. M. I. Love, W. Huber, S. Anders, Moderated estimation of fold change and dispersion for RNA-seq data with DESeq2. Genome Biol 15, 550 (2014).

